# Molecular responses of agroinfiltrated *Nicotiana benthamiana* leaves expressing suppressor of silencing P19 and influenza virus-like particles

**DOI:** 10.1101/2023.07.21.550062

**Authors:** Louis-Philippe Hamel, Rachel Tardif, Francis Poirier-Gravel, Asieh Rasoolizadeh, Chantal Brosseau, Geneviève Giroux, Jean-François Lucier, Marie-Claire Goulet, Adam Barrada, Élise Roussel, Marc-André Comeau, Pierre-Olivier Lavoie, Peter Moffett, Dominique Michaud, Marc-André D’Aoust

**Affiliations:** Medicago Inc., 2552, boul. du Parc-Technologique, Québec, QC, G1P 4S6, Canada; Centre SÈVE, Faculté des Sciences, Département de Biologie, Université de Sherbrooke, Sherbrooke, QC, J1K 2R1, Canada; Centre de recherche et d’innovation sur les végétaux, Département de phytologie, Université Laval, Québec, QC, G1V 0A6, Canada

**Keywords:** Plant immunity, Influenza hemagglutinin, Virus-like particles, Suppressor of RNA silencing P19, Transient *Agrobacterium*-mediated expression, *Nicotiana benthamiana*, Plant molecular farming

## Abstract

The production of influenza vaccines in plants is achieved through transient *Agrobacterium*-mediated expression of viral hemagglutinins (HAs). These proteins are produced and matured through the secretory pathway of plant cells, before being trafficked to the plasma membrane where they induce formation of virus-like particles (VLPs). Production of VLPs unavoidably impacts plant cells, as do viral suppressors of RNA silencing (VSRs) that are often co-expressed to increase protein yields. However, little information is available on host molecular responses to these foreign proteins. The present work provides a comprehensive overview of transcriptomic, metabolic, and signaling changes occurring in *Nicotiana benthamiana* leaf cells transiently expressing the VSR P19, or co-expressing P19 and an influenza HA. Our data identifies generic responses to *Agrobacterium*-mediated expression of foreign proteins, including shutdown of chloroplast gene expression, activation of oxidative stress responses, and reinforcement of the plant cell wall through lignification. Our results also indicate that P19 expression promotes salicylic acid (SA) signaling, a process apparently antagonized by co-expression of HA. As the latter induces specific signatures, with effects on lipid metabolism, lipid distribution, and oxylipin signaling, dampening of P19 responses suggests crosstalk between SA and oxylipin pathways. Consistent with the upregulation of oxidative stress-related genes and proteins, we finally show that reduction of oxidative stress damage through exogenous application of ascorbic acid improves plant biomass quality during production of VLPs.

**One-sentence summary:** *Agrobacterium*-mediated expression of influenza virus-like particles induces a unique molecular signature in *Nicotiana benthamiana* leaf cells.

## Introduction

Hemagglutinins (HAs) are trimeric glycoproteins found on the surface of influenza viruses. HAs are essential for infection as they bind to sialic acid receptors located in the plasma membrane (PM) of epithelial cells from the host respiratory system. Vaccination is broadly recognized as one of the most effective methods to prevent influenza infections and limit societal and economic burden associated to seasonal and pandemic strains of the virus (Ortiz de Lejarazu-Leonardo *et al.,* 2021). Host neutralizing antibodies that recognize the receptor-binding domain of HAs block interaction of the virus with host cells and therefore represent a key correlate of protection against influenza. Correspondingly, HA antigens constitute the main target for commercial production of influenza vaccines. Egg-based approaches are still the most widely employed to produce influenza vaccines, however these methods have remained mostly unchanged since their introduction more 70 years ago (Bouvier, 2018). Egg-based vaccine production is also constrained by certain drawbacks, including year-to-year (or strain-to-strain) variation in effectiveness, and dependency on pathogen-free egg supplies (Rajaram *et al.,* 2020), which may become an issue in case of avian influenza pandemics.

Transient *Agrobacterium tumefaciens*-mediated expression (agroinfiltration) is a powerful tool to express recombinant proteins in plants and is widely employed to study protein function or sub-cellular localization *in planta* (Sainsbury and Lomonossoff, 2014). When performed on a large-scale, agroinfiltration can be considered as a *bona fide* ‘molecular farming’ approach, a term collectively referring to strategies in which plant cells are used as factories to produce biopharmaceutical products (Chung *et al.,* 2022). Molecular farming offers alternative approaches to classical protein production systems, and it has become helpful in the fight against global health issues, including new influenza virus strains, and the emergence of new infectious diseases.

Using a molecular farming approach in *N. benthamiana* leaf cells, the biopharmaceutical company Medicago has developed a method to produce influenza vaccines via transient expression of recombinant influenza *HA* genes (Landry *et al.,* 2010). Engineered to more efficiently enter the secretory pathway of plant cells, newly synthesized HAs are trafficked to the PM, where they end up in discrete microdomains called lipid rafts. When sufficient HA proteins have accumulated, PM curvature is altered, resulting in budding of so-called virus-like particles (VLPs; D’Aoust *et al.,* 2008). These nanoscale assemblies comprise trimer clusters of the engineered HA protein, as well as a lipid envelope derived from the plant cell PM. Structurally, VLPs and influenza virions are similar in size and shape, however the former lack genetic components required for replication. Once purified and formulated into vaccine candidates, VLPs activate an immune response protecting newly immunized hosts from subsequent infection by the virus (Landry *et al.,* 2010).

To sustain commercial production of influenza vaccines in plants, transient expression of HAs was optimized so high quantities of VLPs are produced. This includes co-expression of the viral suppressor of RNA silencing (VSR) P19 (Silhavy *et al.,* 2002), which prevents silencing of *HA* genes delivered by the *Agrobacterium*. Since plant cells produce high amounts of HA proteins and supply membrane lipids to support the formation of VLP envelopes, this expression system unavoidably triggers energy and resource demanding processes, including elevated transcription and translation rates, as well as plant immunity. While the effects of disarmed *Agrobacterium* strains were previously investigated (Ditt *et al.,* 2006; Anand *et al.,* 2008), the combined impacts of agroinfiltration and foreign protein expression, particularly when producing VLPs, have not despite widespread use of *Agrobacterium* to produce recombinant proteins in plants (Goulet *et al.,* 2012; Grosse-Holz *et al.,* 2018; Jutras *et al.,* 2020).

In this study, we combined several approaches to shed light on the complex interplay of responses taking place within plant cells transiently expressing P19 only, or co-expressing P19 and the HA protein from pandemic avian influenza virus strain Indonesia (H5^Indo^), which leads to the formation of VLPs *in planta*. Our results show that *Agrobacterium*-mediated expression of these foreign proteins results in downregulation of chloroplast-related genes (CRGs). Transient expression of the foreign proteins also resulted in the activation of generic defense responses, including lignification and upregulation of genes involved in oxidative stress and systemic acquired resistance (SAR). Importantly, activation levels of commonly induced pathways were higher upon HA protein expression. In contrast, the activation of other molecular responses was more specific to the recombinant protein expressed, including salicylic acid (SA) responses and increased expression of cytosolic heat shock proteins (HSPs) by P19 expression. Likewise, VLP expression resulted in the modulation of lipid metabolism, of lipid distribution within membranes, and in the activation of oxylipin responses. Our results thus show that VLP production in *N. benthamiana* triggers a unique molecular signature that affects plant cells through enhanced expression of genes commonly involved in wounding or herbivory stresses. They also suggest that plant cells producing VLPs experience signal crosstalk, with SA-dependent responses induced by P19 at least partially antagonized by HA expression, perhaps due to oxylipin-related signaling generated by the latter. Better understanding of plant responses to foreign protein expression will help to improve molecular farming techniques, as exemplified here with the exogenous application of ascorbic acid (AsA), an antioxidant that inhibited activation of plant immune responses and improved biomass quality during VLP expression.

## Results

### Stress symptoms and HA protein expression

The effects of *Agrobacterium* infiltration, P19 expression, and production of VLPs in *N. benthamiana* were characterized using multiple approaches. These first included macroscopic evaluation of stress symptoms on representative leaves of each condition harvested 6 days post-infiltration (DPI). Using non-infiltrated (NI) leaves as a baseline, no obvious effect was visible on leaves infiltrated only with buffer (Mock), or infiltrated with *Agrobacterium* strain AGL1 carrying a binary vector control (AGL1; Figure 1A). For AGL1-infiltrated leaves only expressing P19, yellowish discoloration of the leaf blade was seen, suggesting some level of chlorosis. No evidence of plant cell death was denoted on P19 leaves. In comparison, H5 leaves co-expressing P19 and H5^Indo^ showed more pronounced chlorosis, as well as greyish necrotic flecking suggesting early stage of plant cell death activation (Figure 1A). To confirm accumulation of recombinant protein H5^Indo^, a western blot analysis was performed using an antibody specific to the HA protein from influenza virus strain Indonesia. Results showed that H5^Indo^ was only detected in H5 samples (Figure 1B). Correspondingly, hemagglutination (HMG) assays indicated that HA activity was only measurable in samples expressing H5^Indo^ (Figure 1C). HA activity also confirmed that recombinant H5^Indo^ proteins synthesized *in planta* were active against receptors located on red blood cells. Overall, molecular analyses confirmed that collected plant biomass was suitable for downstream analyses, including evaluation of the effects of foreign protein expression on the plant transcriptome.

**Figure 1.**
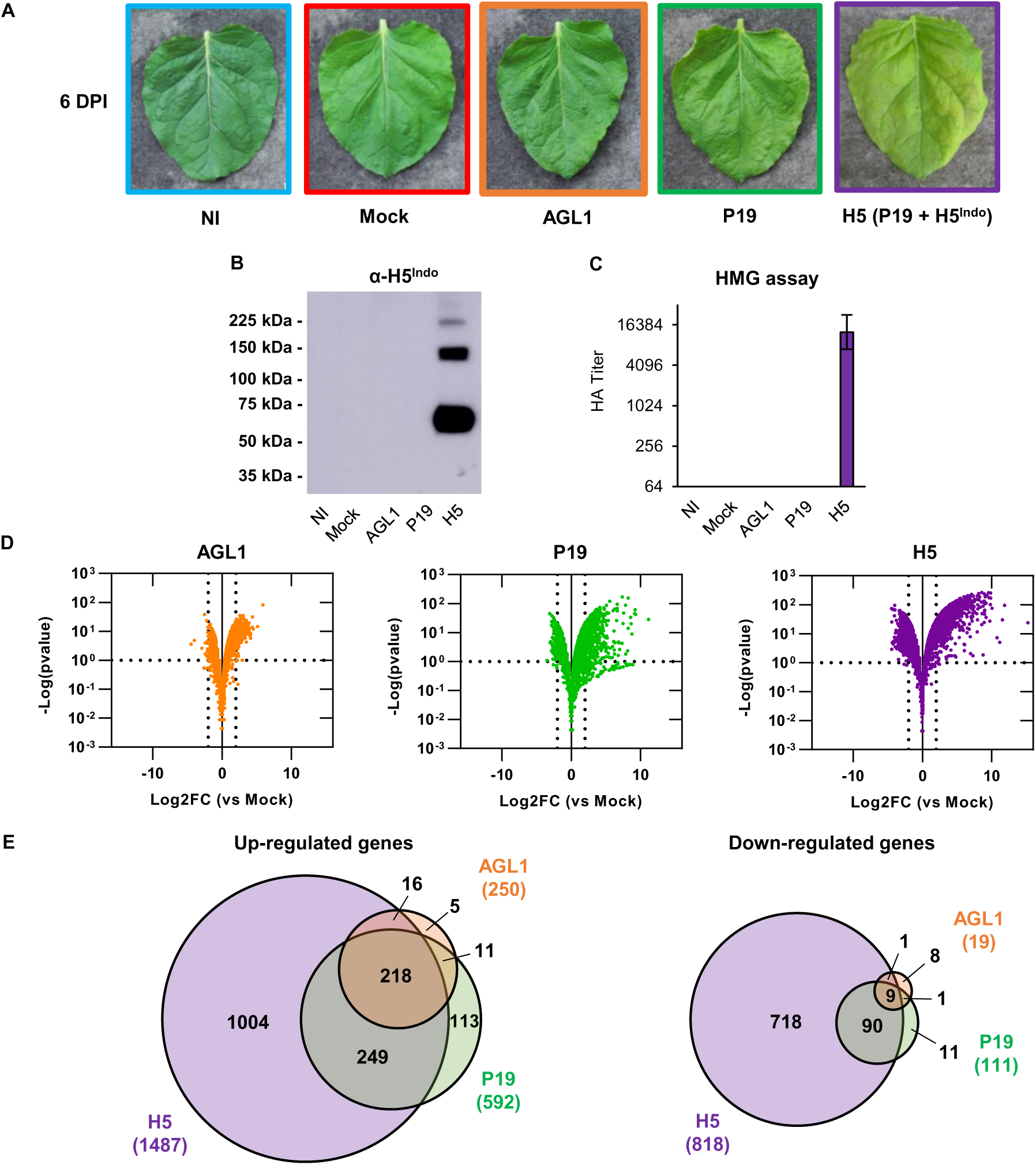
Stress symptoms and analysis of biomass used for transcriptome analyses. (A) Stress symptoms observed on representative leaves from each condition harvested at 6 DPI. (B) Western blot confirming HA protein accumulation. (C) Hemagglutination (HMG) assay confirming HA protein activity Global transcriptional changes at 6 DPI following AGL1 infiltration (left panel), P19 expression (middle panel), or P19 and HA co-expression (right panel), as depicted by volcano plots. Dashed lines represent expression thresholds: Log2FC ≥ 2 or ≤ -2 and padj < 0.1. (E) Venn diagrams depicting up- and downregulated genes from pairwise comparisons: AGL1 *vs* Mock, P19 *vs* Mock, and H5 *vs* Mock. Circle size is proportional to the number of genes significantly regulated. Genes specific to AGL1 infiltration are shown in orange. Genes specific to P19 expression are shown in green. Genes specific to P19 and HA co-expression are shown in purple. Diagram intersects show genes common to more than one condition. Condition names are as follow: NI: non-infiltrated leaves; Mock: leaves infiltrated with buffer only; AGL1: leaves infiltrated with *Agrobacterium* strain AGL1 that carry a binary vector control; P19: leaves infiltrated with AGL1 and expressing P19 only; H5: leaves infiltrated with AGL1 and co-expressing P19 and H5^Indo^.

### Effects of foreign protein expression on the N. benthamiana leaf transcriptome

An RNAseq analysis was conducted with RNA from the tissues described above. Using the Mock treatment as a control (impact of the infiltration without *Agrobacterium*), pairwise comparisons were performed. To be considered significantly regulated, genes had to fulfill the following criteria: Log2 of the expression fold change value (Log2FC) ≥ 2 or ≤ -2 and adjusted p-value (padj) < 0.1 (corresponding to a false discovery rate below 10%). Based on these parameters, volcano plots of all sequenced genes (Figure 1D) and Venn diagrams of up- and downregulated genes (Figure 1E) were created. For each section of Venn diagrams, the unsorted list of up- and downregulated genes is available in the Table S1. At 6 DPI, results showed that AGL1 infiltration resulted in the upregulation of 250 genes and in the downregulation of 19 genes (Figure 1E). These numbers were lower than those observed following expression of P19 only, or co-expression of P19 and H5^Indo^ (Figures 1D and 1E). Venn diagrams also revealed that upregulated genes specific to AGL1 only accounted for 2% of all upregulated genes in this condition (5/250). For downregulated genes, AGL1-specific genes accounted for 42% of the total (8/19), however this number was still low considering the small size of this gene subset (Figure 1E). In other words, gene sets from AGL1 samples largely overlapped with those of the P19 and H5 samples. In light of this, the Mock treatment was kept as control, but focus was placed on P19 and H5 samples for subsequent RNAseq comparisons and confirmation of gene regulation through real time quantitative polymerase chain reaction (RTqPCR).

When comparing P19 and H5 samples, RNAseq revealed that co-expression of the two proteins resulted in up- or downregulation of more genes than the expression of P19 only (Figures 1D and 1E). In addition, most genes with altered expression were specific to H5 samples, suggesting that HA expression induces a unique molecular signature in transformed plant cells. For gene sets that overlapped between P19 and H5 samples, extent of the gene up- or downregulation was also generally higher in HA-expressing samples (Table S1). Globally, transcriptional changes were consistent with intensity of the stress symptoms, which were also increased for H5 samples (Figure 1A).

### Downregulation of chloroplast-related genes

Gene annotation suggested that several nuclear genes encoding for chloroplast-localized proteins were repressed in P19 and H5 samples (Table S1). These included genes encoding chlorophyll-binding or chlorophyll synthesis proteins, protein components of the photosystems, and small subunits of the ribulose-1,5-bisphosphate carboxylase-oxygenase (RuBisCO; Table S2). For H5 samples, the downregulated gene list also comprised gene model Niben101Scf06721g00011, which is annotated as a □Response regulator 18□ (Table S1). Actually, this gene encodes a Golden 2-Like (GLK) protein, a class of transcription factors (TFs) that promotes expression of CRGs (Waters *et al.,* 2009). Using complete gene lists from P19 and H5 pairwise comparisons, all annotated CRGs were extracted regardless of their Log2FC or padj values (Table S2). The volcano plot created using this gene subset revealed that more CRGs were repressed in P19 and H5 samples, although not meeting the minimum Log2FC threshold of -2 (Figure 2A). The volcano plot also highlighted that for most CRGs repressed, the extent of the downregulation was higher in H5 samples compared to P19 samples (Figure 2A). The analysis also revealed two additional *GLK* genes that were repressed in P19 and H5 samples, although these did not meet the minimum Log2FC threshold initially established (Table S2). Also annotated as □Response regulators□, the downregulation of these genes suggests that CRG expression was broadly repressed by foreign protein expression. Indeed, the small number of downregulated genes in AGL1 samples (Figure 1E; Table S1) suggests that at 6 DPI, it is recombinant protein expression rather than agroinfiltration that results in downregulation of CRGs.

**Figure 2.**
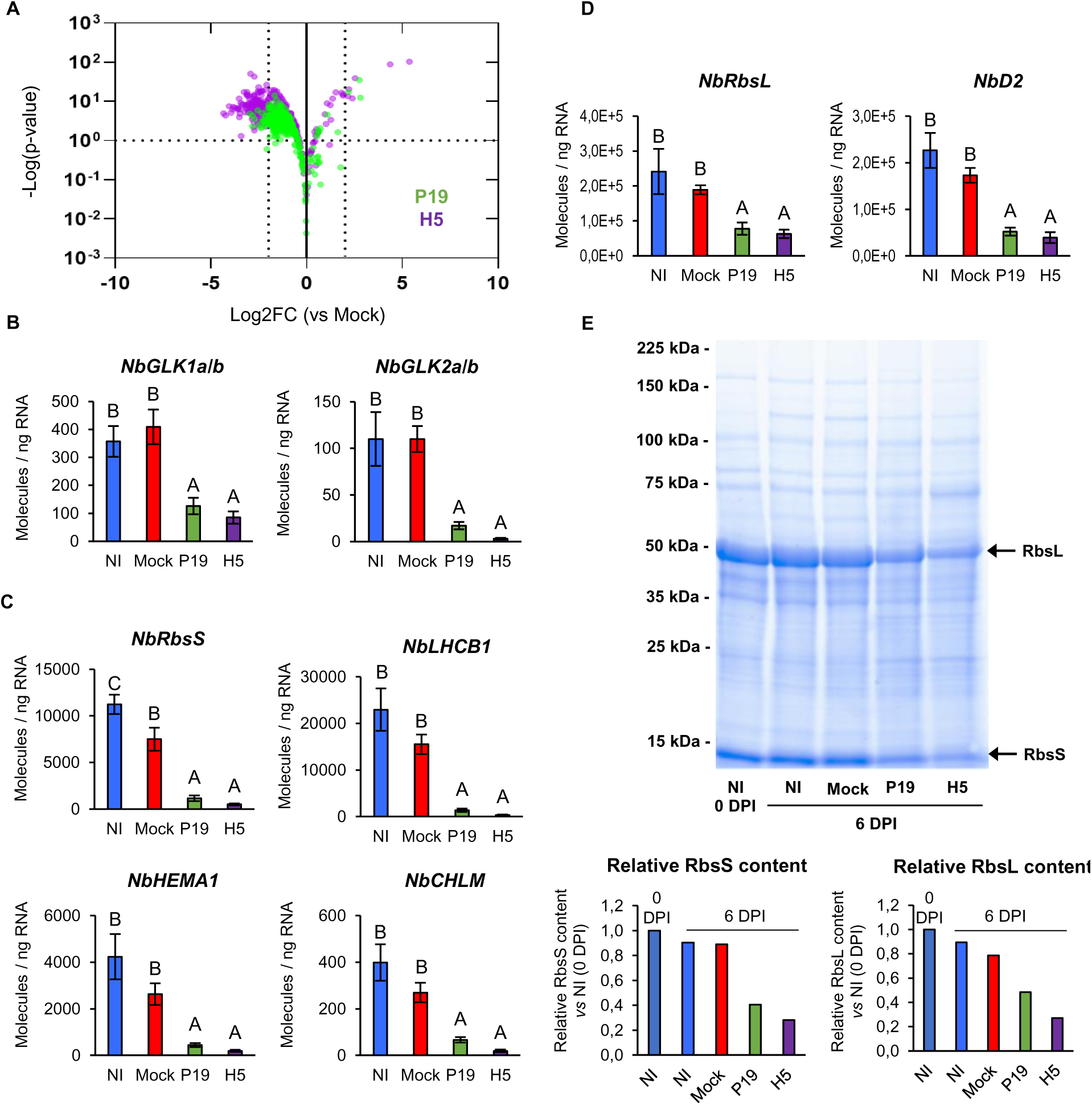
Downregulation of chloroplast-related genes and decreased RuBisCO content. (A) Volcano plot depicting impact of P19 expression (green) or P19 and HA co-expression (purple) on CRGs at 6 DPI. Comparisons of whole transcriptomics data were made using the Mock treatment as a control. Dashed lines represent expression thresholds: Log2FC ≥ 2 or ≤ -2 and padj < 0.1. Expression of *NbGLK* genes (B), of nuclear genes encoding for chloroplast-localized proteins (C), and of genes located on the chloroplast genome (D) as measured by RTqPCR at 6 DPI. Results are expressed in numbers of molecules per ng of RNA. Groups that do not share the same letter are statistically different. (D) Total protein extracts from various conditions harvested at 0 or 6 DPI. Following SDS-PAGE analysis, gel was stained with Coomassie blue. Arrows highlight RuBisCO small and large subunits (RbsS and RbsL, respectively). Underneath bar graphs depict relative RbsS and RbsL content as measured by densitometry. Rbs subunit content from leaves harvested prior to infiltration (0 DPI) were arbitrarily set at one-fold. Condition names are as follow: NI: non-infiltrated leaves; Mock: leaves infiltrated with buffer only; P19: leaves infiltrated with AGL1 and expressing P19 only; H5: leaves infiltrated with AGL1 and co-expressing P19 and H5^Indo^.

To confirm downregulation of CRGs, RTqPCR was performed. Tested candidates included two pairs of closely related *GLKs* (Figure 2B), as well as nuclear genes encoding chloroplast-localized proteins (Figure 2C). Primers specific to genes located on the chloroplast genome were also developed (Figure 2D). In all cases examined, RTqPCR confirmed similar or slightly lower transcript levels when comparing NI and Mock samples. In contrast, expression of CRGs was significantly reduced in P19 and H5 samples compared to NI and Mock controls. Further indication of altered chloroplast function came from the analysis of total protein extracts visualized after Coomassie blue staining (Figure 2E). At 6 DPI, NI and Mock samples displayed similar levels of both RuBisCO subunits. These levels were also similar to those observed in NI samples harvested prior to infiltration (0 DPI). For P19 and H5 samples at 6 DPI, levels of both RuBisCO subunits had clearly decreased compared to NI and Mock controls.

### Upregulation of HSP and chaperone genes following P19 expression

RNAseq revealed that 113 genes were specifically upregulated in P19 samples (Figure 1E). Among these, 36 (∼32%) encode cytosolic HSPs or other types of molecular chaperones (Tables S1 and S3). Using whole gene lists from P19 and H5 pairwise comparisons, *HSP* and chaperone genes were extracted regardless of their Log2FC or padj values (Table S3). The volcano plot created using this gene subset highlighted specificity of the response in P19 samples, despite co-expression of the VSR in H5 samples (Figure 3A). As expression of only three *HSPs* was slightly induced in AGL1 samples (Table S1), results suggest that it is P19 expression that really drives this response. To confirm this induction pattern, primers targeting the highly induced *HSP* genes Niben101Scf04040g09011 (*NbHSP1a*) and Niben101Scf10306g00024 (*NbHSP1b*), as well as Niben101Scf04490g00001 (*NbHSP70*) and Niben101Scf03114g03011 (*NbHSP90*) were designed. In all cases, RTqPCR confirmed a strong upregulation of *HSP* genes in P19 samples compared to other conditions (Figure 3B).

**Figure 3.**
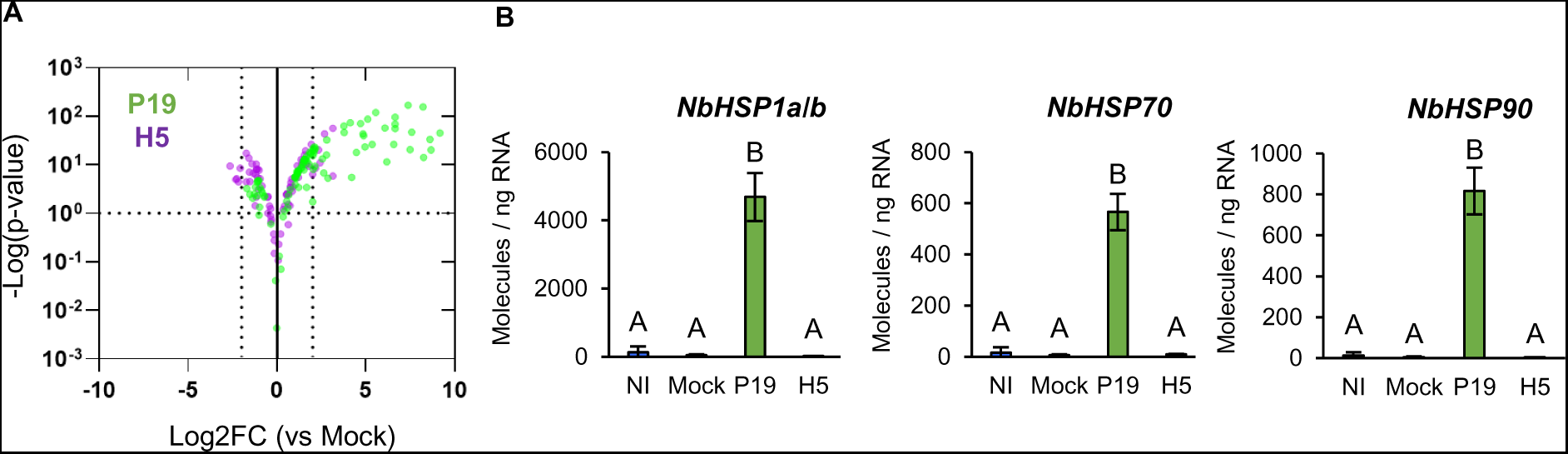
Upregulation of *HSP* and chaperone genes by P19 expression. (A) Volcano plot depicting impact of P19 expression (green) or P19 and HA co-expression (purple) on *HSP* and chaperone genes at 6 DPI. Comparisons of whole transcriptomics data were made using the Mock treatment as a control. Dashed lines represent expression thresholds: Log2FC ≥ 1 or ≤ -1 and padj < 0.1. (B) Expression of *HSP* genes as measured by RTqPCR. Results are expressed in numbers of molecules per ng of RNA. Groups that do not share the same letter are statistically different. Condition names are as follow: NI: non-infiltrated leaves; Mock: leaves infiltrated with buffer only; P19: leaves infiltrated with AGL1 and expressing P19 only; H5: leaves infiltrated with AGL1 and co-expressing P19 and H5^Indo^.

### Upregulation of lipid-related genes

Secretion of the HA protein requires the endomembrane system of plant cells, while the lipid envelope of VLPs is derived from the PM of the same plant cells (D’Aoust *et al.,* 2008). Consistent with this, RNAseq identified several lipid-related genes, including some that encode two closely related phospholipases D (PLDs), one phospholipase C (PLC), and two diacylglycerol kinases (DGKs; Tables S1 and S4). In all cases, genes were induced in P19 samples, however upregulation levels were higher in H5 samples. While PLDs convert structural phospholipids into phosphatidic acid (PA), PLCs produce soluble inositol 1,4,5-trisphosphate and diacyl glycerol (DAG) that remains in membranes, but can be converted to PA by DGKs (Canonne *et al.,* 2011). The *PLD* genes identified were Niben101Scf02465g00004 (*NbPLD*β*1*) and Niben101Scf16022g04010 (*NbPLD*β*2*), which are most closely related to *PLD*β*1* of *Arabidopsis thaliana*. The latter promotes pathogen-induced production of JA, which in turn represses SA signaling (Zhao *et al.,* 2013). *PLC* gene Identified was Niben101Scf02221g00009 (*NbPLC2*), which is most closely related to Arabidopsis *PLC2*, whose protein product promotes the production of reactive oxygen species (ROS) during immunity (D’ Ambrosio *et al.,* 2017). The most highly induced *DGK* gene was Niben101Scf06654g02004 (*NbDGK5*), a close homolog of *NtDGK5* that promotes oxidative stress in tobacco (Cacas *et al.,* 2017). To confirm RNAseq results, RTqPCR was carried out with primers specific to *NbPLDs*, *NbPLC2,* and *NbDGK5*. In all cases, results confirmed slight upregulation in P19 samples, and a significantly higher gene induction in H5 samples (Figure 4A). RNAseq also revealed strong and H5-specific upregulation of the Niben101Scf10067g02017 gene (Tables S1 and S4), which encodes a putative glycerol-3-phosphate acyltransferase (GPAT). This class of enzymes converts glycerol-3-phosphate into lysophosphatidic acid (LPA). RTqPCR confirmed that this gene, herein termed *NbGPAT5*, was highly induced in a manner dependent on HA protein expression (Figure 4A). Taken together, our results suggests that HA protein expression favors the accumulation of PA and LPA, changes that could be due to an increased demand for certain lipid species associated to HA protein secretion, and/or VLP budding.

**Figure 4.**
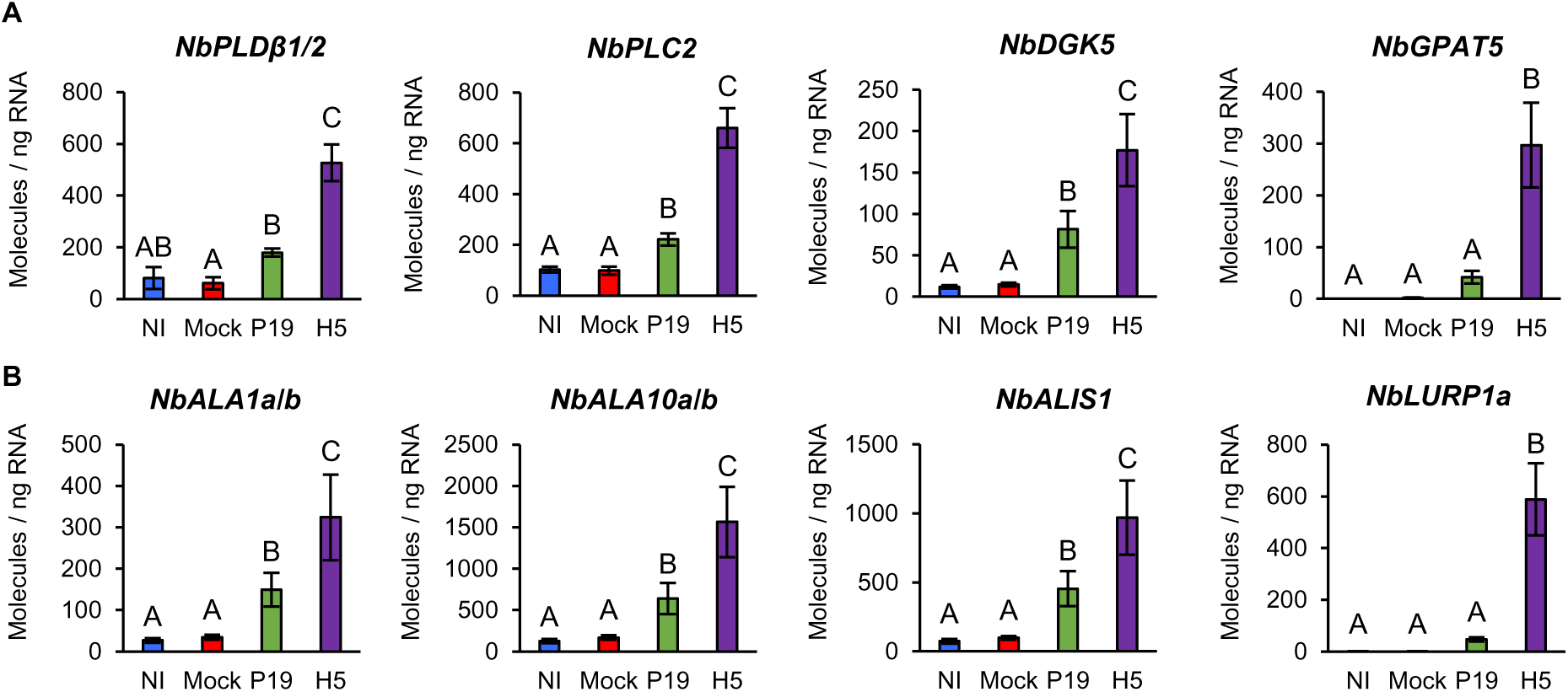
Expression of lipid-related genes. Expression of genes involved in lipid metabolism and signaling (A), or in distribution of lipids within membranes (B) as measured by RTqPCR at 6 DPI. Results are expressed in numbers of molecules per ng of RNA. Groups that do not share the same letter are statistically different. Condition names are as follow: NI: non-infiltrated leaves; Mock: leaves infiltrated with buffer only; P19: leaves infiltrated with AGL1 and expressing P19 only; H5: leaves infiltrated with AGL1 and co-expressing P19 and H5^Indo^.

Under normal conditions, the PM is characterized by an asymmetric distribution of its lipid components and the alteration of this asymmetry affects PM curvature (van Meer *et al.,* 2008). Modulation of lipid asymmetry is thus an important feature for vesicle-mediated secretion, in addition of being reported during host-virus interactions and activation of plant cell death (López-Marqués *et al.,* 2014). In mammalian cells, lipid asymmetry is controlled by proteins that flip lipids from one leaflet to the other, including flippases and scramblases (Devaux *et al.,* 2006). In Arabidopsis, P4 ATPases known as ALAs have been shown to work as flippases (López-Marqués *et al.,* 2014), while proteins of the DUF567 family have structural features reminiscent of scramblases (Bateman *et al.,* 2009). RNAseq data revealed that *ALA* homologs (*NbALAs*) were induced in P19 samples, but that upregulation was higher in H5 samples (Tables S1 and S4). Among induced *NbALAs* were close relatives of Arabidopsis *ALA10* that internalizes exogenous phospholipids across the PM (Poulsen *et al.,* 2015). Also induced were homologs of *ALA3* and *ALA1*, which encode for proteins that localize to the Golgi apparatus and PM, respectively (Poulsen *et al.,* 2008; López-Marqués et al., 2012). These ALAs also interact with ALA-Interacting Subunits (ALISs), which contribute to their sub-cellular localization and activity. A homolog of Arabidopsis *ALIS1* was also induced in P19 and H5 samples, with a higher upregulation level again observed in the latter (Tables S1 and S4). For Niben101Scf02167g00009 (*NbALA1a*) and Niben101Scf21069g00002 (*NbALA1b*), as well as Niben101Scf03029g01010 (*NbALA10a*) and Niben101Scf00465g02013 (*NbALA10b*), RTqPCR confirmed significantly higher expression in H5 samples compared to P19 samples (Figure 4B). A similar expression pattern was also observed for Niben101Scf02639g03015 (*NbALIS1*).

Interestingly, RNAseq also revealed strong, and in most cases H5-specific, induction of genes homologous to Arabidopsis *late upregulated in response to Hyaloperonospora parasitica 1* (*LURP1*; Tables S1 and S4). While required for immunity mediated by resistance proteins (Knoth and Eulgem, 2008), *LURP1* encodes a protein of the DUF567 family and thus possess structural features reminiscent of lipid scramblases. For Niben101Scf00819g08011 (*NbLURP1a*), RTqPCR confirmed strong and H5-specific gene upregulation (Figure 4B). Again, these transcriptional changes may be needed to alter membrane composition and architecture during HA secretion and/or VLP budding.

### Upregulation of oxidative stress-related genes

In plant cells, membrane respiratory burst oxidase homologs (RBOHs) are a major source of ROS (Møller *et al.,* 2007), with Arabidopsis RBOHd and RBOHf reported to be involved in plant immunity (Torres *et al.,* 2002). At 6 DPI, RNAseq showed that Niben101Scf02581g04013 (*NbRBOHd*) and Niben101Scf10840g01010 (*NbRBOHf*) were induced in a manner largely dependent on HA protein expression (Tables S1 and S5). For *NbRBOHd*, this observation was confirmed by RTqPCR (Figure 5A). ROS are also produced by electron transport chains from stressed mitochondria and chloroplasts, as well as via the activity of cellular oxidases (van Aken and van Breusegem, 2015). Among the most highly induced genes specific to H5 samples, RNAseq identified two closely related *polyphenol oxidases* (*PPOs*; Tables S1 and S5). Typically activated in response to wounding and herbivory, PPOs oxidize phenolic compounds that in turn react with oxygen and proteins to form ROS (Tran *et al.,* 2012). RTqPCR with primers specific to Niben101Scf00180g08002 (*NbPPO1*) and Niben101Scf04384g02014 (*NbPPO3*) confirmed HA protein expression to result in strong and specific upregulation of these genes (Figure 5B).

**Figure 5.**
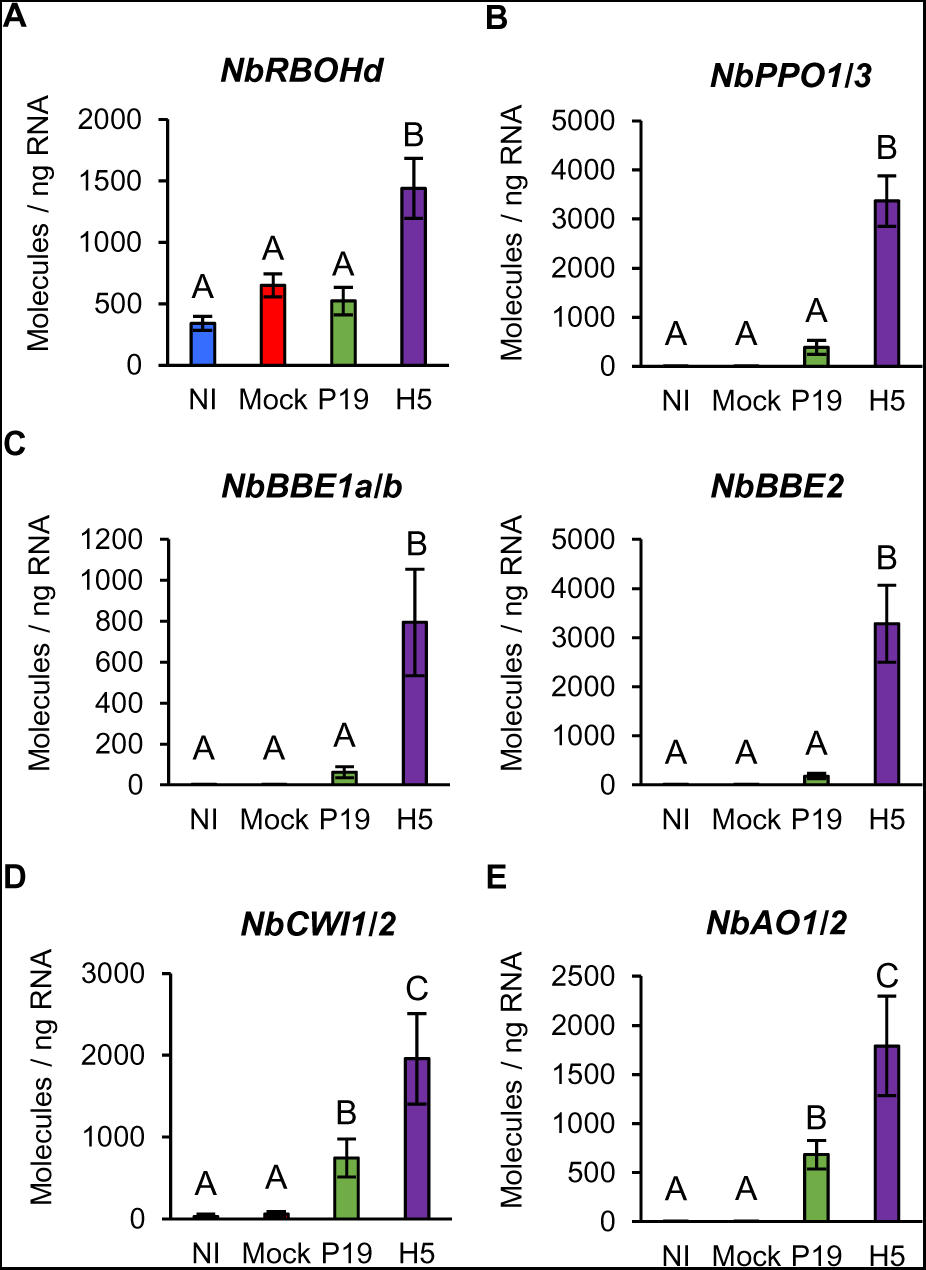
Expression of genes linked to the activation of oxidative stress. Expression of genes encoding for NADPH oxidases (A), PPOs (B), secreted carbohydrate oxidases of the BBE family (C), CWIs (D), or secreted AOs (E) as measured by RTqPCR at 6 DPI. Results are expressed in numbers of molecules per ng of RNA. Groups that do not share the same letter are statistically different. Condition names are as follow: NI: non-infiltrated leaves; Mock: leaves infiltrated with buffer only; P19: leaves infiltrated with AGL1 and expressing P19 only; H5: leaves infiltrated with AGL1 and co-expressing P19 and H5^Indo^.

In response to HA expression, several genes encoding for berberine-bridge enzymes (BBEs) were also among the most highly induced (Tables S1 and S5). BBEs belong to a functionally diverse protein family, however these oxidases all produce hydrogen peroxide (Daniel *et al.,* 2017). BBEs have been involved in a number of cellular processes, however the most highly induced *NbBBE* genes all encoded secreted proteins sharing homology to carbohydrate oxidases (Carter and Thornburg, 2004; Custers *et al.,* 2004). This suggests that HA protein expression promotes sugar oxidation in the apoplast, which may in turn contribute to the activation of oxidative stress. In line with this, RNAseq revealed that expression of genes related to sugar metabolism in the apoplast was induced in P19 samples, but even more so in H5 samples (Tables S1 and S6). These included homologs of sugar transport proteins and cell wall invertases (CWIs) that participate in plant defense (Lemonnier *et al.,* 2014; Veillet *et al.,* 2016). For *BBE* genes Niben101Scf01395g03002 (*NbBBE1a*) and Niben101Scf01061g07011 (*NbBBE1b*), as well as Niben101Scf00944g01001 (*NbBBE2*), strong and H5-specific induction was confirmed by RTqPCR (Figure 5C). RTqPCR also confirmed induction of the *CWI* genes Niben101Scf12270g02002 (*NbCWI1*) and Niben101Scf04632g03006 (*NbCWI2*) in both P19 and H5 samples, with again upregulation levels significantly higher in the latter (Figure 5D). Taken together these results suggest that foreign protein expression promotes conversion of sucrose to simple sugars in the apoplast, possibly providing energy to support local activation of defense. However, in HA-expressing samples, increased accumulation of simple sugars in the apoplast may contribute to ROS generation via protein products of *NbBBE* genes that are strongly induced in this condition (Figure 5B).

To prevent extensive damage caused by ROS, plant cells produce antioxidant metabolites, including ascorbic acid (AsA) that is the major antioxidant of the apoplast (Pignocchi *et al.,* 2006). RNAseq showed that closely related *ascorbate oxidase* (*AO*) genes were upregulated in AGL1, P19, and H5 samples, with a higher upregulation level in the latter (Tables S1 and S5). As the redox status of apoplastic AsA is negatively controlled by secreted AOs (Pignocchi and Foyer, 2003), our results suggest that upregulation of *NbAO* genes contributes to oxidative stress activation in the apoplast, especially when the HA protein is expressed. RTqPCR with primers specific to Niben101Scf03026g01009 (*NbAO1*) and Niben101Scf22432g00001 (*NbAO2*) confirmed upregulation of these genes in P19 and H5 samples compared to controls, with again higher expression level in the latter (Figure 5E).

### Lignin-related gene expression and lignin quantification

In response to pathogens, plant cells reinforce their cell wall through the coordinated deposition of polymers, including lignin (Wang *et al.,* 2013). The lignin biosynthesis pathway comprises numerous classes of enzymes that catalyse synthesis and cell wall polymerisation of monolignol precursors (Figure 6A). When expressing secreted VLPs, lignification poses extra challenges to extract and purify desired product from the reinforced plant cell wall. RNAseq suggested specific or at least stronger expression of several genes with predicted function in lignification (Tables S1 and S7). Notably, secreted *peroxidase* (*PRX*) and *laccase* (*LAC*) genes were among the most highly induced in response to HA protein expression. Also upregulated were monolignol synthesis genes, including *4-coumarate-CoA ligases* (*4CLs*), a *hydroxycinnamoyl-CoA transferase* (*HCT*), and *caffeoyl-CoA O-methyltransferases* (*CCoAOMTs*; Table S7). When displayed as a heat map, our results showed that for H5 samples, most steps of the lignin biosynthesis pathway were represented by at least one upregulated gene (Figure 6B). To confirm these results, RTqPCR was performed on selected genes either involved in monolignol synthesis (Figure 6C), or lignin polymerization (Figure 6D). Compared to controls, results showed that genes were slightly induced by expression of P19 only, but that co-expression of P19 and HA proteins resulted in significantly higher gene induction. To confirm lignin accumulation during foreign protein expression, lignin content was quantified in NI, P19, and H5 samples harvested at 6 DPI (Figure 6E). These results showed that P19 and H5 samples accumulated more lignin than the NI control, again with a much stronger effect in H5 samples.

**Figure 6.**
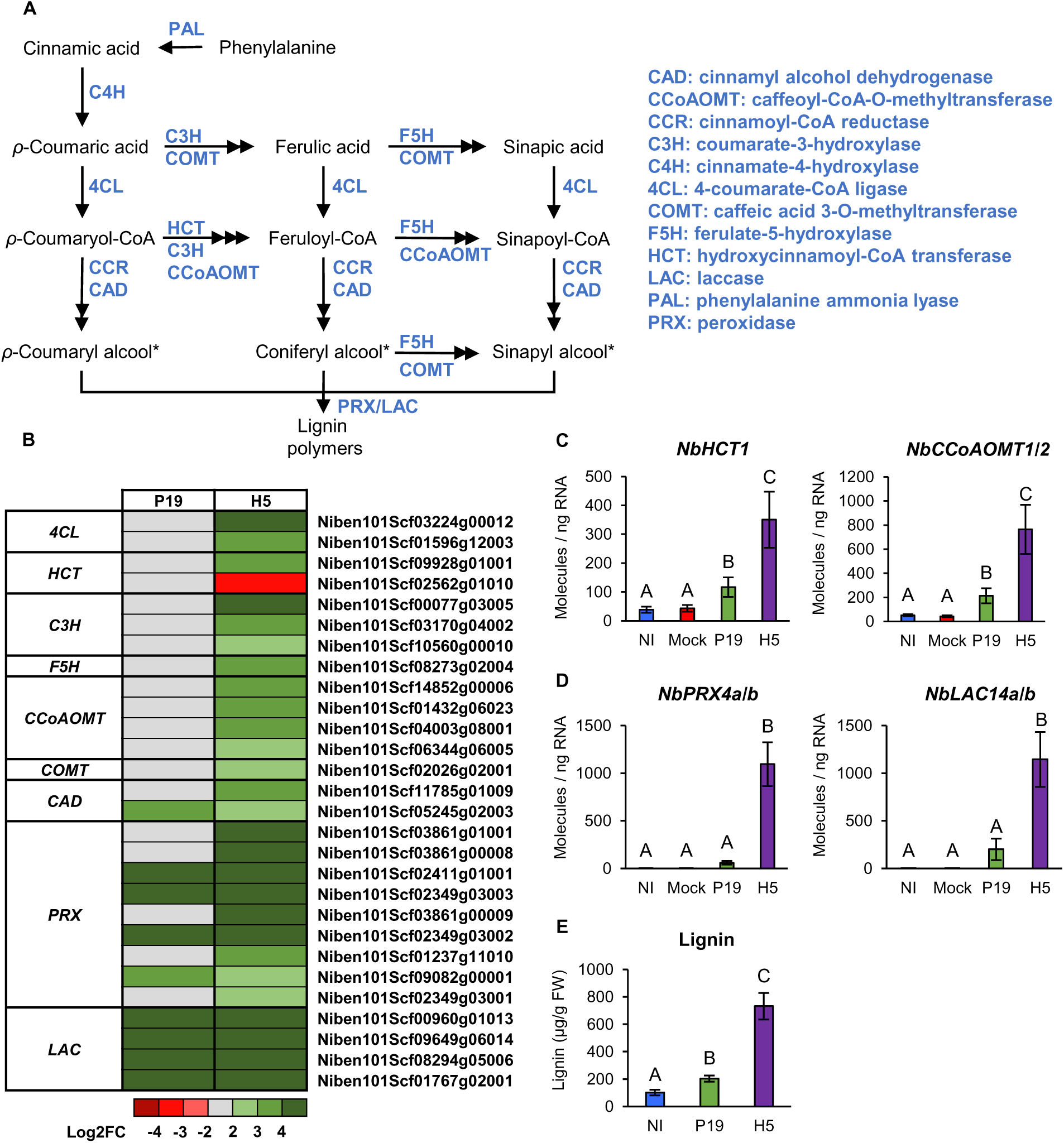
Expression of lignin-related genes and lignin quantification. (A) Overview of the lignin synthesis pathway. Metabolites are shown in black, while enzymes are shown in blue. Monolignol precursors of lignin are marked with an asterisk. (B) Heat-map depicting expression of genes involved in monolignol precursor synthesis or lignin polymerization in the cell wall at 6 DPI. Each line represents a gene shown in the Table S7. Grey indicates genes that are not differentially expressed. Green are red colored gradients respectively reflect extent of gene up- and downregulation, as indicated. At 6 DPI, RTqPCR confirms upregulation of monolignol precursor synthesis genes (C) as well as genes involved in lignin polymerization (D). Results are expressed in numbers of molecules per ng of RNA. (E) Lignin accumulation at 6 DPI. Results are expressed in ug of lignin per g of biomass fresh weight (FW). Groups that do not share the same letter are statistically different. Condition names are as follow: NI: non-infiltrated leaves; Mock: leaves infiltrated with buffer only; P19: leaves infiltrated with AGL1 and expressing P19 only; H5: leaves infiltrated with AGL1 and co-expressing P19 and H5^Indo^.

### Salicylic acid synthesis and signaling

To shed light on stress hormone signaling during foreign protein expression, SA was quantified in NI, AGL1, P19, and H5 samples. At 6 DPI, NI samples showed barely detectable levels of SA, while significantly higher SA accumulation was observed in P19 samples compared to AGL1 and H5 samples (Figure 7A). From these results, we deduced that AGL1 infiltration induces moderate accumulation of SA and that expression of P19 increases this response. In H5 samples, which co-express P19 and HA proteins, P19-mediated accumulation of SA appeared to be compromised, perhaps because HA-induced signaling interferes with this process. Alternatively, P19 levels may be lower when the VSR is co-expressed with the HA protein. In AGL1, P19 and H5 samples, most salicylates were found in a conjugated form, with highest levels detected in H5 samples compared to P19 and even more so AGL1 samples (Figure 7A).

**Figure 7.**
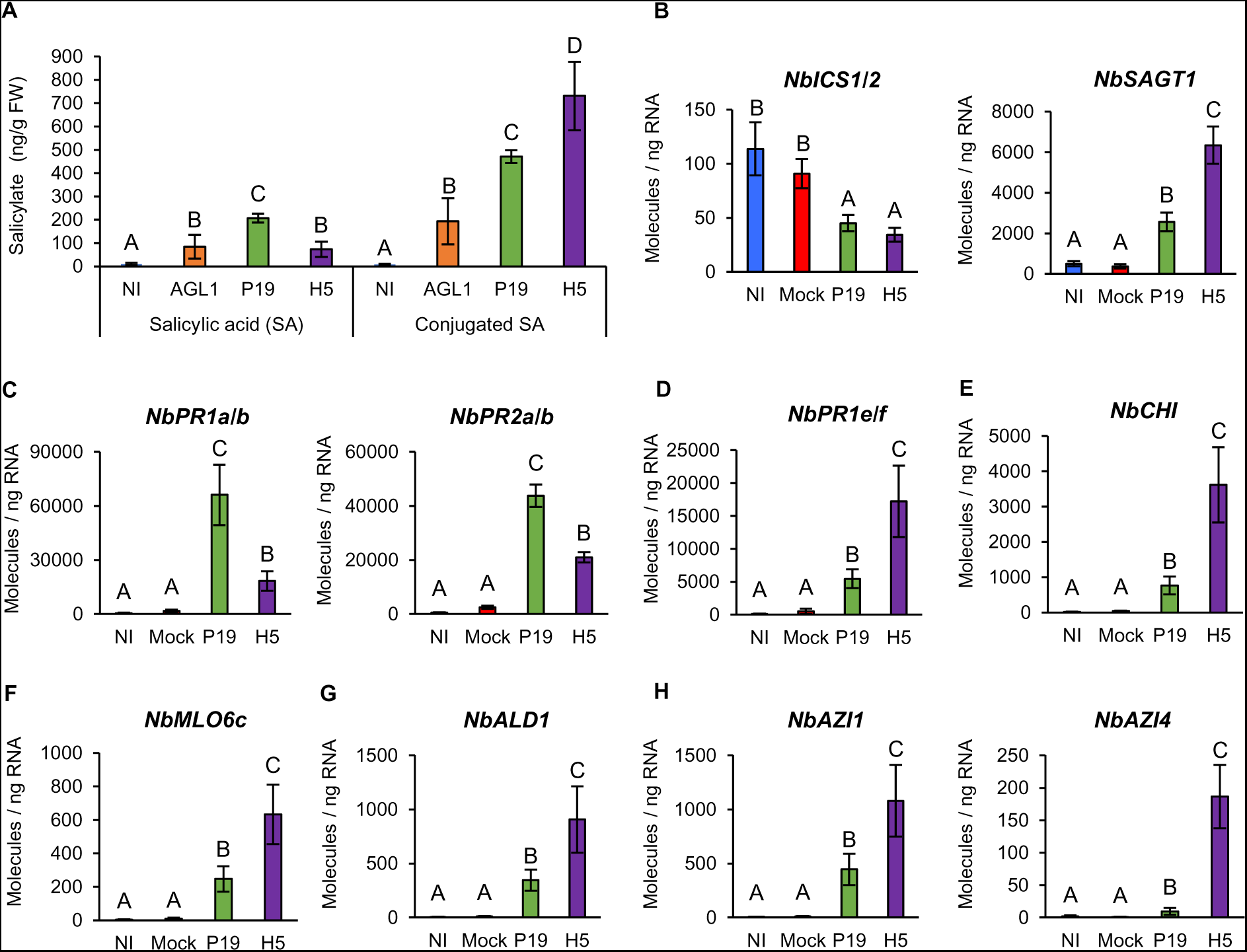
Salicylate accumulation and expression of SA- and SAR-related genes. (A) Accumulation of SA and conjugated SA at 6 DPI. Results are expressed in ng of salicylate per g of biomass fresh weight (FW). RTqPCR confirms that salicylate accumulation is coupled to the expression of SA regulatory genes (B) and of SA response genes (C). RTqPCR was also performed to assess expression of different SAR response genes (D, E, F) and of SAR regulatory genes (G, H). RTqPCR results are expressed in numbers of molecules per ng of RNA. Groups that do not share the same letter are statistically different. Condition names are as follow: NI: non-infiltrated leaves; Mock: leaves infiltrated with buffer only; AGL1: leaves infiltrated with *Agrobacterium* strain AGL1 that carry a binary vector control; P19: leaves infiltrated with AGL1 and expressing P19 only; H5: leaves infiltrated with AGL1 and co-expressing P19 and H5^Indo^.

In plants, SA is produced via two independent pathways (Chen *et al.,* 2009). In *N. benthamiana*, stress-induced production of SA however depends on the expression of *isochorismate synthase* (*ICS*) genes (Catinot *et al.,* 2008). RNAseq indicated that Niben101Scf00593g04010 (herein termed *NbICS1*) was repressed in both P19 and H5 samples (Tables S1 and S8). RTqPCR targeting *NbICS1* and close homolog Niben101Scf05166g06006 (*NbICS2*) confirmed reduced *ICS* gene expression in both P19 and H5 samples (Figure 7B). This suggests that P19-induced accumulation of SA (Figure 7A) is either not dependent on *NbICS* gene upregulation, or that this upregulation occurred before harvesting at 6 DPI. Alternatively, this response may be mediated through the alternative SA biosynthesis route, although *phenylalanine ammonia lyase* (*PAL*) genes, which encode the first enzyme of this pathway, were not induced in the tested conditions.

In Arabidopsis, decreases in SA levels have been associated with transcriptional repression of *ICS1* and increased SA storage caused by upregulation of the *SA glucosyl transferase 1* (*SAGT1*) gene Zheng *et al.,* 2012). Protein product of the latter gene converts SA into inactive conjugated forms that are stored for later use (Dean and Delaney, 2008). In *N. benthamiana*, Niben101Scf05415g00003 is the closest homolog of *SAGT1*. Here termed *NbSAGT1*, RNAseq showed that this gene was induced in P19 samples, but that upregulation level was higher in H5 samples (Tables S1 and S8). This expression pattern was confirmed by RTqPCR (Figure 7B), indicating that transcriptomics correlated with detected levels of conjugated SA (Figure 7A).

To further characterize SA signaling, expression of *Pathogenesis Related 1* (*PR1*) genes was investigated. The genome of *N. benthamiana* comprises at least 20 *PR1-like* genes, with Niben101Scf01999g07002, Niben101Scf00107g03008, Niben101Scf03376g03004, and Niben101Scf13926g01014 being the most similar to *AtPR1* (At2g14610; Figure S1), the Arabidopsis homolog widely employed as a marker of SA signaling. The first three of these genes, respectively termed *NbPR1a*, *NbPR1b*, and *NbPR1c*, were identified in RNAseq data. Expression was higher in P19 samples compared to AGL1 and H5 samples (Tables S1 and S8), a pattern that correlated with SA accumulation levels (Figure 7A). RTqPCR with primers specific to *NbPR1a* and *NbPR1b* confirmed higher induction in P19 samples compared to H5 and furthermore control samples (Figure 7C). Interestingly, similar expression patterns were observed for β*-1,3-glucanase* genes of the *PR2* family, which are also often employed as markers of SA signaling (Durrant and Dong, 2004). In these cases, RNAseq revealed similar induction levels in AGL1 and P19 samples, while expression was lower in H5 samples (Tables S1 and S8). For Niben101Scf01001g00003 (*NbPR2a*) and Niben101Scf01001g00005 (*NbPR2b*), RTqPCR confirmed higher gene upregulation in P19 samples compared to other conditions (Figure 7C).

### Expression of SAR-related genes

Examination of RNAseq data highlighted the upregulation of a second *PR1* gene subset, which interestingly behaved differently compared to their previously described homologs (see above). In this case, Niben101Scf04053g02007 (*NbPR1e*), Niben101Scf04053g02006 (*NbPR1f*), and Niben101Scf00953g03009 (*NbPR1g*) were induced in AGL1 and P19 samples, but expression was more pronounced in H5 samples (Tables S1 and S8). Those three genes are closely related and encode structurally distinct PR1 proteins that possess a short C-terminal extension (CTE) following the conserved CAPE1 peptide of PR1 proteins (Figure S1; Chen et al., 2014). RTqPCR with primers specific to *NbPR1e* and *NbPR1f* confirmed the distinct transcriptional behavior of these *PR1* genes, with highest expression levels detected in H5 samples (Figure 7D).

Enhanced *PR1* gene expression is not only associated to SA, but also to the SAR (Durrant and Dong, 2004). Expression of SAR-related genes was thus examined, including *N. benthamiana* homologs of *PR3* gene *Chitinase* (*CHI*) and *Mildew Resistance Locus O 6 (MLO6*) of Arabidopsis (Riedlmeier *et al.,* 2017). RNAseq revealed that several *PR3* genes were induced in AGL1, P19, and H5 samples, with higher upregulation levels seen in the latter (Tables S1 and S8). This included Niben101Scf02171g00007 (*NbCHI*), the closest homolog of Arabidopsis *CHI*. For *MLO6* homologs, several candidates were specifically induced in H5 samples, while others were induced in both P19 and H5 samples, again with higher transcript levels in the latter (Tables S1 and S8). This was, for instance, the case for Niben101Scf07792g02034 (*NbMLO6c*), the closest homolog of Arabidopsis *MLO6*. As for *NbPR1e* and *NbPR1f* (Figure 7D), RTqPCR confirmed upregulation of *NbCHI* (Figure 7E) and *NbMLO6c* (Figure 7F) in P19 samples compared to controls, however induction was significantly higher in H5 samples.

To further confirm that SAR is part of the response to HA protein expression, RNAseq data was searched for SAR regulatory genes, including homologs of Arabidopsis *AGD2-Like Defense 1* (*ALD1*; *Song et al.,* 2004) and *Azelaic Acid-Induced 1* (*AZI1*; Cecchini *et al.,* 2015). While Niben101Scf04547g02001 (*NbALD1*) was induced in AGL1 and P19 samples, upregulation level was higher in H5 samples (Tables S1 and S8). For *AZI1* homologs, Niben101Scf13429g02004 (*NbAZI1*) and Niben101Scf13429g03011 (*NbAZI5*) were similarly induced in AGL1 and P19 samples, while induction was higher in H5 samples. For Niben101Scf04779g01029 (*NbAZI2*), Niben101Scf09387g02004 (*NbAZI3*), and Niben101Scf07599g00019 (*NbAZI4*), enhanced expression was only detected in H5 samples (Tables S1 and S8). For *NbALD1* (Figure 7G) and *NbAZI* genes (Figure 7H), expression patterns observed by RNAseq were confirmed via RTqPCR, including specific induction of *NbAZI4* in H5 samples. Taken together, these results suggest that at 6 DPI, SAR-related responses were induced in AGL1 and P19 samples, however activation of this pathway was stronger and broader in HA-expressing samples. Distinct expression patterns of *PR1* gene subsets also suggest that PR1 proteins with a CTE are more indicative of SAR, while PR1 proteins without a CTE were as expected indicative of SA signaling.

### Oxylipin synthesis and signaling

As discussed for *PPO* genes (Figure 5B), transcriptomics suggests that wounding and herbivory response genes are induced by HA protein expression. Indeed, a search of RNAseq data revealed H5-specific induction of several genes that typically respond to these stresses, including those that encode for plant defensins (PDFs), serine protease inhibitors (PIs), Kunitz trypsin inhibitors (KTIs), and cysteine PIs of the PR4 family (annotated as □wound-induced proteins□; Tables S1 and S9). For the plant defensin gene Niben101Scf17290g01005 (*NbPDF1*) and *PI* genes Niben101Scf00294g01014 (*NbPI2*), Niben101Scf06424g00003 (*NbKTI3*), and Niben101Scf00773g08003 (*NbPR4b*), RTqPCR confirmed strong and H5-specific upregulation (Figure 8A).

**Figure 8.**
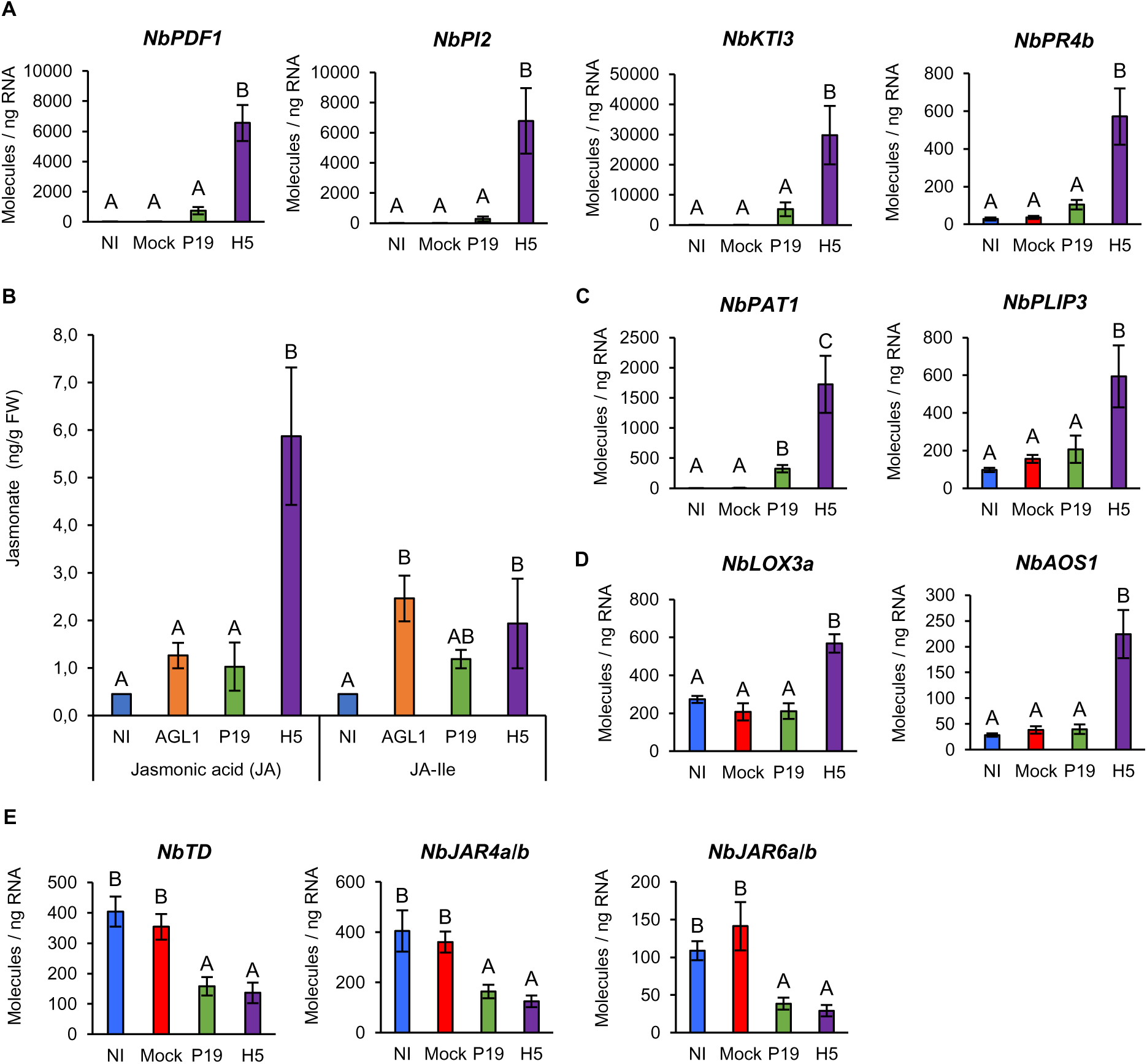
Jasmonate accumulation and expression of oxylipin-related genes. (A) Expression of oxylipin response genes as measured by RTqPCR at 6 DPI. (B) Accumulation of JA and JA-Ile at 6 DPI. Results are expressed in ng of jasmonate per g of biomass FW. RTqPCR was also performed on genes encoding for PLAs (C), as well as on genes involved in JA (D) and JA-Ile (E) biosynthesis. RTqPCR results are expressed in numbers of molecules per ng of RNA. Groups that do not share the same letter are statistically different. Condition names are as follow: NI: non-infiltrated leaves; Mock: leaves infiltrated with buffer only; AGL1: leaves infiltrated with *Agrobacterium* strain AGL1 that carry a binary vector control; P19: leaves infiltrated with AGL1 and expressing P19 only; H5: leaves infiltrated with AGL1 and co-expressing P19 and H5^Indo^.

In plants, wounding and herbivory responses often depend on jasmonic acid (JA; Wasternack and Feussner, 2018). Upregulation of *PDF* and *PI* genes thus prompted evaluation of JA levels in NI, AGL1, P19, and H5 samples. While low JA levels were detected in samples from the first three conditions, significantly higher JA accumulation was detected in H5 samples (Figure 8B). Accordingly, RNAseq revealed upregulation of many *phospholipase A* (*PLA*) genes, in a manner that was largely dependent on HA protein expression (Tables S1 and S9). These included closely related Niben101Scf03309g01011 (*NbPLA1*) and Niben101Scf01228g02012 (*NbPLA2*), which encode secreted proteins. Also induced were several *patatin* (*PAT*) genes encoding proteins closely related to *Nicotiana tabacum* homologs that promote accumulation of oxylipins, including JA (Dhondt *et al.,* 2002; Cacas *et al.,* 2005). Closely related Niben101Scf01076g05026 and Niben101Scf01623g13002, which encode homologs of Arabidopsis PLASTID LIPASEs (AtPLIPs), were also identified. Localized in chloroplasts, PLIPs also promote the accumulation of bioactive oxylipins in response to stress (Wang *et al.,* 2018). RTqPCR with primers specific to Niben101Scf06277g00007 (*NbPAT1*) and Niben101Scf01623g13002 (*NbPLIP3*) confirmed their upregulation, with patterns mostly specific to HA protein expression (Figure 8C).

To synthesize JA, free fatty acids produced by PLAs are oxygenated by 13-lipoxygenases (13-LOXs), before cyclization by allene oxide synthases (AOSs) and allene oxide cyclases (AOCs). Resulting intermediate 12-oxophytodienoic acid (OPDA) then translocates to peroxisomes where reductases and β-oxidation cycles complete JA synthesis (Wasternack and Feussner, 2018). Consistent with JA accumulation in H5 samples (Figure 8B), *13-LOX* gene Niben101Scf02749g01001 (*NbLOX3a*), *AOS* gene Niben101Scf05799g02010 (*NbAOS1*), and *OPDA reductase* (*OPR*) gene Niben101Scf00779g06009 (*NbOPR1*) were all specifically upregulated in H5 samples (Tables S1 and S9). For *NbLOX3a* and *NbAOS1*, RTqPCR confirmed H5-specific upregulation of gene expression (Figure 8D).

To become entirely bioactive, JA is conjugated to isoleucine (Ile; Staswick and Tiryaki, 2004). Quantification of jasmonate-isoleucine (JA-Ile) revealed overall low levels of the compound, with non-significant differences observed between NI and P19 samples, and a small but significant increase in AGL1 and H5 samples relative to NI samples but not P19 samples (Figure 8B). In tobacco, herbivore-induced formation of JA-Ile requires the *Jasmonate-Resistant 4* and *6* (*JAR4/6*) genes (Wang *et al.,* 2008), as well as the gene *threonine deaminase* (*TD*) that is involved in biosynthesis of Ile (Kang *et al.,* 2006). At the threshold examined, *N. benthamiana* homologs from these genes were not identified by RNAseq as being differentially expressed. However, RTqPCR analysis revealed that Niben101Scf02502g14001 (*NbTD*) was repressed in P19 and H5 samples compared to NI and Mock controls (Figure 8E). Similarly, RTqPCR analyses with primers targeting closely related Niben101Scf05584g01007 (*NbJAR4a*) and Niben101Scf22940g00002 (*NbJAR4b*), as well as Niben101Scf00470g00001 (*NbJAR6a*) and Niben101Scf01083g00008 (*NbJAR4b*) showed significantly reduced expression in P19 and H5 samples compared to controls (Figure 8E). These results were consistent with overall low accumulation of JA-Ile (Figure 8B).

Strong H5-specific upregulation of wounding and herbivory response genes, coupled to low JA-Ile accumulation, suggest production of other bioactive oxylipins in response to HA protein expression. Aside from JA and JA-Ile, the octadecanoid pathway allows for the synthesis of other oxylipins with roles in defense (Figure 9A; Wasternack and Feussner, 2018). To shed light on which of these metabolites may be active in H5 samples, RNAseq data was searched for more oxylipin regulatory genes (Figure 9B). This revealed several candidates involved in alternative branches of the octadecanoid pathway. All identified genes were induced in a manner specific or at least more pronounced in H5 samples (Tables S1 and S9). These included *9-lipoxygenase* (*9-LOX*) genes, among which Niben101Scf01434g03006 (*NbLOX1*) was the most highly induced (Figure 9B). Also identified was Niben101Scf04626g00009, an α*-dioxygenase* (*DOX*) gene termed *NbDOX1*. In Arabidopsis, closest homologs of *NbLOX1* and *NbDOX1* work in conjugation to promote synthesis of oxylipins that activate local and systemic defenses (Vicente *et al.,* 2012). A number of cytochromes P450 of family 74 (CYP74s) were also identified (Figure 9B; Tables S1 and S9). Improperly annotated as *AOS* genes, these candidates actually encode divinyl ether synthases (DESs) or epoxyalcohol synthases (EASs) that are related to, but functionally distinct from AOSs that promote JA synthesis (Figure 9A). In tobacco, NtDES1 works in coordination with 9-LOX enzymes to produce divenyl ether fatty acids involved in defense (Fammartino *et al.,* 2007). Our analyses also identified Niben101Scf05133g06002 (*NbEH1*) and Niben101Scf00640g04023 (*NbEH2*), closely related *epoxide hydrolase* (*EH*) genes upregulated in AGL1 and P19 samples, but even more so in H5 samples (Figure 9B; Tables S1 and S9). Often associated to cell detoxification, EHs also act downstream of EASs to produce oxylipin diols with signaling and anti-microbial functions (Figure 9A; Morisseau, 2013). RTqPCR analyses on *NbLOX1*, *NbDOX1*, and *NbCYP74* genes confirmed increased expression in P19 and H5 samples compared to controls. In all cases, upregulation levels were however significantly higher in H5 samples (Figure 9C). Taken as a whole, our data suggests that VLP expression results in stronger activation of the octadecanoid pathway compared to agroinfiltration, or expression of P19 only. In addition to JA, oxylipins produced by alternative branches of the octadecanoid pathway appear to be actively signaling, especially in HA-expressing samples.

**Figure 9.**
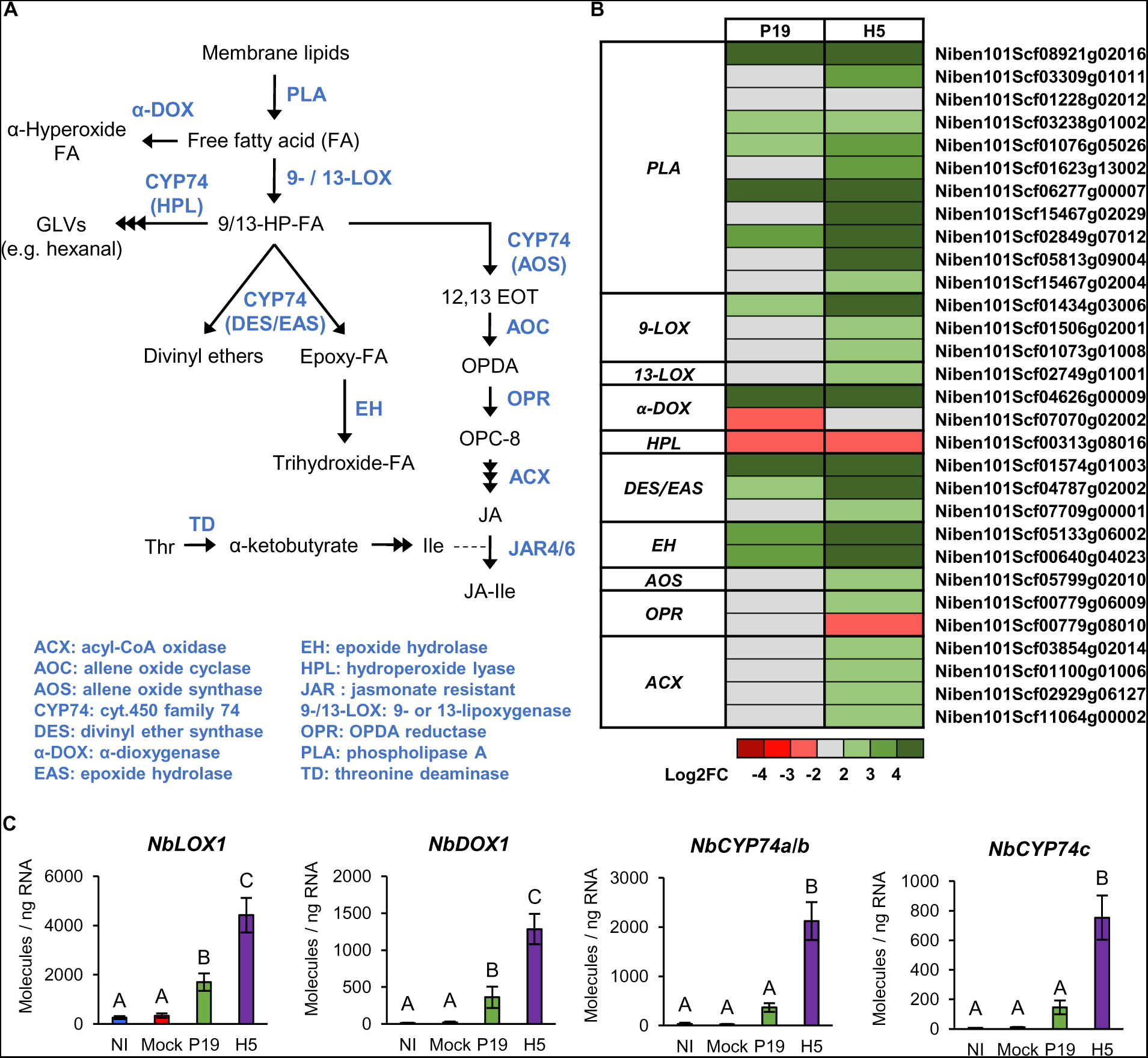
Expression of additional oxylipin regulatory genes. (A) Overview of the octadecanoid pathway. Metabolic intermediates are shown in black while enzymes are shown in blue. (B) Heat-map depicting expression of genes involved in oxylipin metabolism at 6 DPI. Each line represents a gene shown in the Table S9. Grey indicates genes that are not differentially expressed. Green are red colored gradients respectively reflect extent of gene up- and downregulation, as indicated. (C) Expression of genes involved in the synthesis of oxylipins other than JA as measured by RTqPCR at 6 DPI. Results are expressed in numbers of molecules per ng of RNA. Groups that do not share the same letter are statistically different. Condition names are as follow: NI: non-infiltrated leaves; Mock: leaves infiltrated with buffer only; P19: leaves infiltrated with AGL1 and expressing P19 only; H5: leaves infiltrated with AGL1 and co-expressing P19 and H5^Indo^.

### Ethylene and senescence-related pathways

Along with JA, ethylene (ET) is involved in resistance to necrotrophic pathogens, acting antagonistically with SA (Binder, 2020). To assess ET signaling as a result of foreign protein expression, levels of ET precursor 1-aminocyclopropane-1-carboxylate (ACC) were evaluated in NI, AGL1, P19, and H5 samples. At 6 DPI, ACC was not detected in NI samples, while AGL1 and P19 samples displayed low and not significantly different levels of ACC (Figure 10A). In contrast, H5 samples displayed significantly higher level of ACC, suggesting that ET signaling is part of the molecular response to HA protein expression. Accordingly, RNAseq revealed upregulation of several ET biosynthesis genes, including *ACC synthases* (*ACSs*) and *ACC oxidases* (*ACOs*). While some genes were similarly induced in AGL1, P19 and H5 samples, upregulation of other candidates was more pronounced or even specific to HA expression (Tables S1 and S10). For *ACS* gene Niben101Scf09512g03008 (*NbACS3*) and *ACO* gene Niben101Scf08039g01005 (*NbACO4*), RTqPCR confirmed significantly higher upregulation in H5 samples compared to other conditions (Figure 10B).

**Figure 10.**
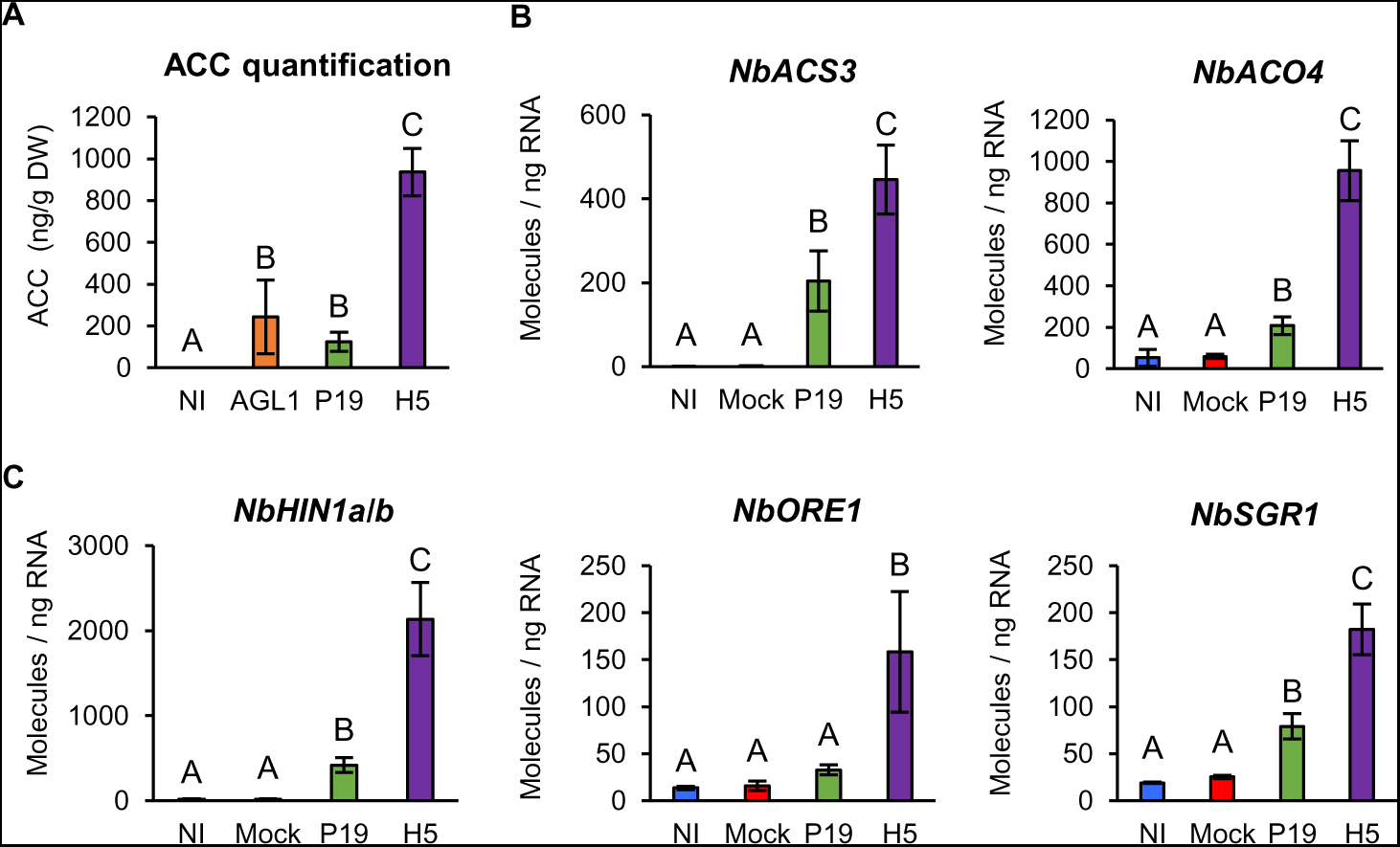
ACC accumulation and expression of ET- or senescence-related genes. (A) Accumulation of ET precursor ACC at 6 DPI. Results are expressed in ng of ACC per g of biomass dry weight (DW). RTqPCR confirms that ACC accumulation is coupled to the expression of genes involved in ET synthesis (B). RTqPCR was also performed on senescence-related genes (C). RTqPCR results are expressed in numbers of molecules per ng of RNA. Groups that do not share the same letter are statistically different. Condition names are as follow: NI: non-infiltrated leaves; Mock: leaves infiltrated with buffer only; AGL1: leaves infiltrated with *Agrobacterium* strain AGL1 that carry a binary vector control; P19: leaves infiltrated with AGL1 and expressing P19 only; H5: leaves infiltrated with AGL1 and co-expressing P19 and H5^Indo^.

ET is also associated with senescence, a form of programmed cell death linked to plant development (Binder, 2020). At 6 DPI, RNAseq indicated that several genes associated with leaf senescence were induced AGL1 and P19 samples, but that upregulation levels were higher in H5 samples (Tables S1 and S10). These included homologs of tobacco asparagine synthetases, which are involved in nitrogen remobilization during senescence (Bovet *et al.,* 2019). Also identified were homologs of the tobacco gene *harpin-induced 1* (*NtHIN1*), which is induced during hypersensitive response and senescence (Takahashi *et al.,* 2004). Close homologs of Arabidopsis genes *ORESARA 1* (*ORE1*) and *STAYGREEN 1* (*SGR1*) were also identified, and these were specifically induced in H5 samples (Tables S1 and S10). While ORE1 stimulates senescence by antagonizing GLK TFs (Rauf *et al.,* 2013), SGR1 promotes senescence-related degradation of chlorophyll (Sakuraba *et al.,* 2015). For Niben101Scf08020g06001 (*NbHIN1a*) and Niben101Scf04717g02003 (*NbHIN1b*), as well as Niben101Scf01277g00002 (*NbORE1*) and Niben101Scf00490g01040 (*NbSGR1*), RTqPCR confirmed higher or even specific gene upregulation in H5 samples (Figure 10C).

### Large-scale proteomics

To investigate changes in protein abundance, new sets of Mock, AGL1, P19 and H5 samples were harvested at 6 DPI. After evaluation of leaf symptoms (Figure S2A), HA protein accumulation and activity were confirmed by western blotting and HMG assays, respectively (Figures S2B and S2C). Using total protein extracts, a proteomics analysis was then performed using the isobaric tags for relative and absolute quantitation (iTRAQ) method. Using the Mock treatment as control, pairwise comparisons were performed for AGL1, P19, and H5 samples. To be considered significantly changed in abundance, proteins had to display a Log2FC value ≥ 1 or ≤ -1. From generated protein lists, Venn diagrams of up- and downregulated proteins were created (Figure S2D). For each diagram section, the unsorted list of up- or downregulated proteins, along with their deduced Z-score, is available in the Table S11.

At the threshold examined, 194 proteins were upregulated, including 89 (∼46%) common to AGL1, P19, and H5 samples (Figure S2D). On the opposite, 75 proteins were downregulated, including 22 (∼30%) common to all conditions. In line with the transcriptome, AGL1 infiltration altered the abundance of fewer proteins compared to P19 expression. In turn, expression of P19 only altered the abundance of fewer proteins compared to P19 and HA co-expression. Many proteins with altered abundance also turned out to be specific to H5 samples, again highlighting induction of a unique molecular signature upon HA expression. As for the transcriptome, abundance of proteins shared between H5 samples and other conditions (AGL1, P19, or both) was generally more affected when the HA protein was expressed (Table S11).

Out of the 16 proteins specifically upregulated in P19 samples (Figure S2D), half were cytosolic HSPs (Table S11). Corresponding genes had all been identified in RNAseq data (Table S3). Several PIs were also upregulated, including NbPI2 in AGL1 and H5 samples, as well as NbKTI1, NbKTI3, and NbPR4b in P19 and H5 samples (Table S11). In all cases, protein accumulation was at least two times higher in H5 samples compared to AGL1 or P19 samples. Genes encoding identified PIs had also been identified in RNAseq data (Table S9). Enhanced accumulation of oxylipin synthesis enzymes was also observed, including NbOPR1 and NbOPR3, that were specific to H5 samples (Table S11). While Niben101Scf00779g06009 (*NbOPR1*) had been identified in the transcriptome (Figure 9B), Niben101Scf05804g03006 (*NbOPR3*) was not, at least at the threshold examined. Oxylipin-related proteins NbLOX1 and NbEH1 were also among upregulated proteins identified, and both accumulated to higher levels in H5 samples compared to AGL1 or P19 samples (Table S11). Again, the corresponding genes had been identified in the transcriptome (Table S9).

Among upregulated proteins, secreted enzymes linked to activation of oxidative stress in the apoplast were also identified. These were putative carbohydrate oxidase NbBBE2 and lignin-forming peroxidase NbPRX4a, which were both specific to H5 samples (Table S11). Also identified were apoplastic sugar invertases NbCWI1 and NbCWI2, as well as ascorbate oxidase NbAO2. These proteins were identified in AGL1, P19, and H5 samples, however with greater accumulation levels in the latter (Table S11). For all secreted enzymes, protein accumulation pattern matched with the expression profile of the corresponding genes in the transcriptome.

Consistent with activation of plant immunity, upregulation of PR proteins was also detected, including SA markers NbPR2a and NbPR2b. In agreement with SA accumulation levels (Figure 7A) and expression of corresponding *PR2* genes (Figure 7C), these proteins accumulated to higher levels in AGL1 and P19 samples compared to H5 samples (Table S11). Again, this supports the idea that SA-mediated signaling was induced by agroinfiltration and furthermore by P19 expression, but that this response was partially antagonized in H5 samples. Peptides belonging to NbPR1e and NbPR1f, which harbor a CTE (Figure S1), were also identified by proteomics. In both cases, protein accumulation was higher in H5 samples compared to AGL1 or P19 samples (Table S11). This pattern was also observed for NbCHI, a marker of SAR. As expression profiles from *NbPR1e*, *NbPR1f*, and *NbCHI* (Figures 7D and 7E) matched protein accumulation patterns, our results support the idea that PR1 homologs with a CTE better reflect activation of SAR and that this pathway is part of the plant response to VLP expression.

### Proteins linked to the unfolded protein response

As described above, proteomics data generally correlated with results from the transcriptome. Interestingly, proteomics also highlighted accumulation of proteins generally involved in the unfolded protein response (UPR). This pathway allows cells to cope with increasing needs in protein secretion, in addition of preventing deleterious effects provoked by accumulation of misfolded proteins in the endoplasmic reticulum (ER; Duwi Fanata *et al.,* 2013). Agroinfiltration coupled to enforced expression of a complex secreted protein such as H5^Indo^ would, for instance, be expected to cause ER stress and to activate the UPR. Accordingly, proteomics identified protein disulfide isomerases (PDIs) with ER retention signals, as well as ER-resident chaperones of the calreticulin (CRT) and binding immunoglobulin protein (BiP) families (proteins marked with an asterisk in the Table S11). Intriguingly, the corresponding UPR genes were not identified in the transcriptome at 6 DPI. This suggests that accumulation of UPR proteins was independent of transcriptional regulation, or that upregulation of UPR genes occurred earlier and in a transient fashion.

### Ascorbic acid reduces HA-induced defenses and stress symptoms

Several lines of evidence suggested that *Agrobacterium*-mediated expression of influenza HA results in oxidative stress activation, especially in the apoplast where VLP accumulation takes place (D’Aoust *et al.,* 2008). We hypothesized that this response was responsible for necrotic symptoms observed on HA-expressing leaves (Figure 1A). To test this, infiltrated plants co-expressing P19 and H5^Indo^ were sprayed with a 10 mM solution of AsA. In *N. benthamiana*, sodium ascorbate was previously shown to suppress necrosis induced by expression of human proteins (Nosaki *et al.,* 2021). To prevent interference with AGL1 infection and T-DNA transfer, the first spray was applied at 2 DPI. Recall treatments were then performed at 48 h intervals, before harvesting at 7 DPI. At harvest, no symptom was observed on NI leaves, while leaves only expressing P19 showed chlorosis (Figure 11A). For leaves co-expressing P19 and H5^Indo^, cell death symptoms of similar intensities were observed for untreated plants, or plants sprayed with a Mock solution. For HA-expressing leaves sprayed with the AsA solution, yellowish discoloration was observed, but little to no necrotic symptoms were denoted. Exogenous application of the antioxidant thus reduced stress symptoms otherwise associated to HA protein expression.

**Figure 11.**
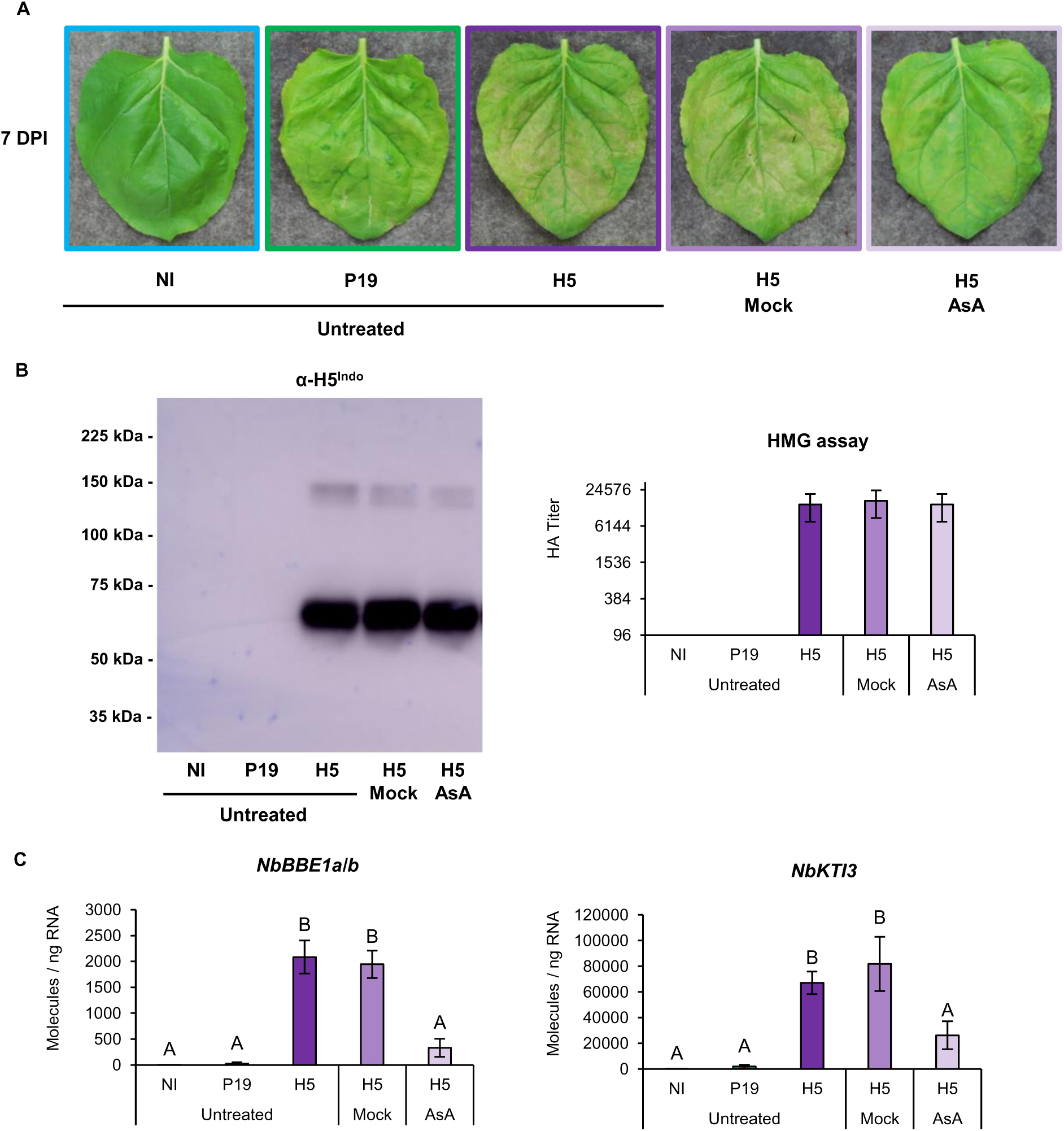
Ascorbic acid reduces HA-induced defenses and symptoms. (A) Stress symptoms observed on representative leaves from each condition harvested at 7 DPI. (B) Western blot confirming HA protein accumulation (left panel). Hemagglutination (HMG) assay confirming HA protein activity (right panel). (C) Expression of HA-induced genes *NbBBE1a/b* and *NbKTI3* as measured by RTqPCR. Results are expressed in numbers of molecules per ng of RNA. Groups that do not share the same letter are statistically different. Condition names are as follow: NI: non-infiltrated leaves; P19: leaves infiltrated with AGL1 and expressing P19 only; H5: leaves infiltrated with AGL1 and co-expressing P19 and H5^Indo^; H5 Mock: HA-expressing leaves sprayed at 48 h intervals with a Mock solution; H5 AsA: HA-expressing leaves sprayed at 48 h intervals with a 10 mM solution of ascorbic acid (AsA).

To confirm that decreased symptoms were not simply caused by lower accumulation of H5^Indo^, HA accumulation and activity were monitored by western blotting and HMG assays, respectively. Results showed similar HA protein levels and similar HA activity in untreated, Mock-treated, or AsA-treated leaves expressing H5^Indo^ (Figure 11B). This confirmed that AsA improved biomass quality at harvest without interfering with HA protein expression nor activity. To assess whether AsA also reduced defense signaling associated with HA protein expression, RTqPCR was performed using genes previously characterized as good markers of HA-induced oxidative stress (*NbBBE1a/b*; Figure 5C), or oxylipin response (*NbKTI3*; Figure 8A). In both cases, results showed barely detectable transcript levels in NI and P19 samples (Figure 11C), confirming the HA-specific regulation of these genes. For untreated and Mock-treated leaves expressing H5^Indo^, genes were similarly induced. Gene upregulation was also detected in H5 leaves sprayed with AsA, however expression levels were significantly lower (Figure 11C), in line with the reduced intensity of necrotic symptoms (Figure 11A).

## Discussion

In characterizing the effects of *Agrobacterium*-mediated expression of foreign proteins in *N. benthamiana*, we found a strong convergence in the datasets collected through RNAseq, RTqPCR, and proteomics. These gene and protein expression patterns were also highly consistent with quantification of defense metabolites, including lignin and stress hormones. Taken as a whole, these results thus provide a comprehensive overview of the complex interplay of responses taking place following expression of P19, or co-expression of this VSR with a VLP-forming protein (Figure 12). Transient expression of these foreign proteins pointed to a trade-off between plant growth and immunity, as exemplified by the downregulation of CRGs and the concomitant activation of many defense pathways. Among the latter, some were co-induced by AGL1 infiltration, expression of P19 only, or co-expression of P19 and H5^Indo^. Generally, co-induced pathways were however induced at higher levels when the HA protein was expressed. On the other hand, some defense responses appeared to be more specific to a defined condition, including enhanced expression of HSPs in P19 samples, or induction of lipid- and oxylipin-related responses in H5 samples (Figure 12).

**Figure 12.**
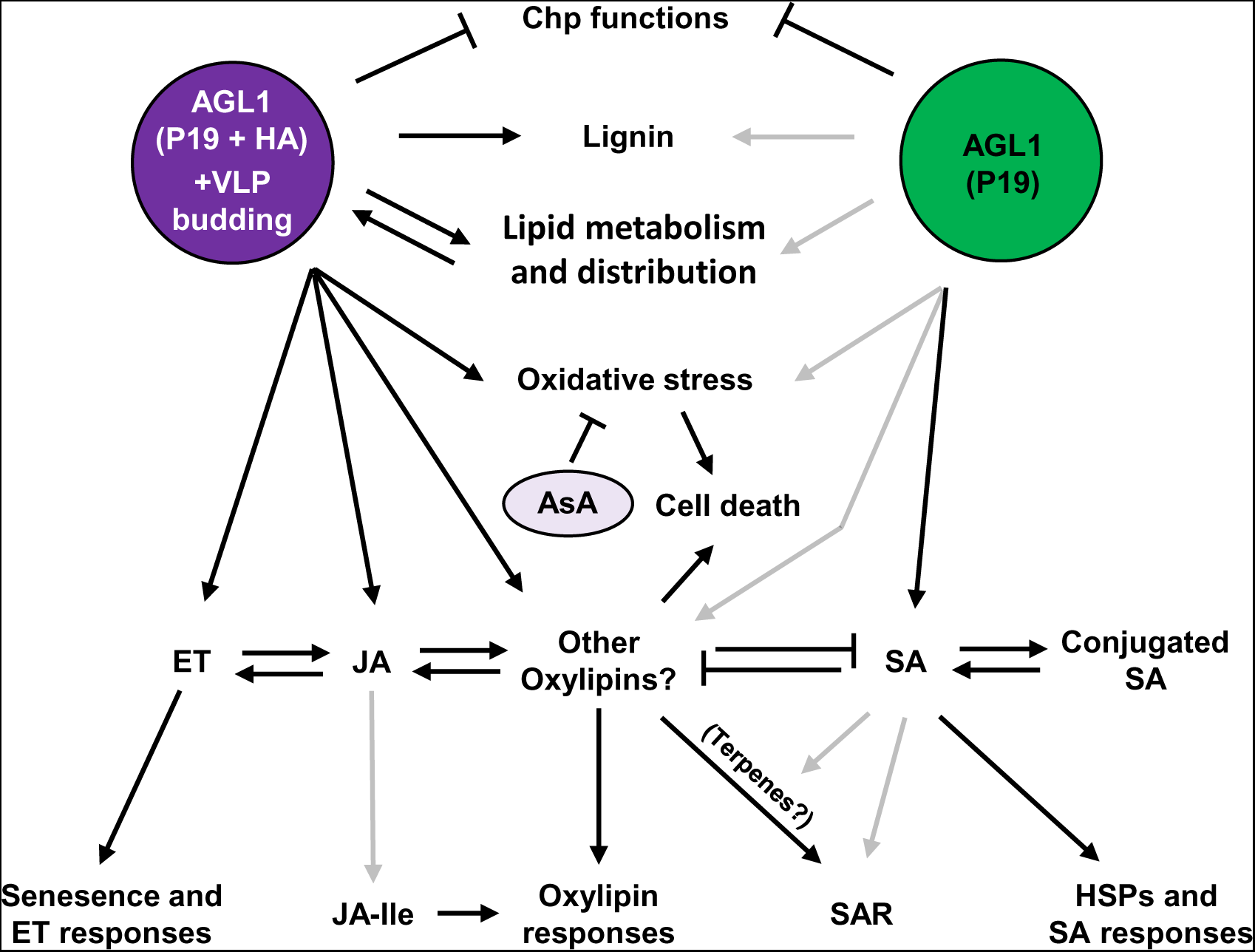
Model of plant responses to P19 and VLP expression. In response to foreign protein expression, some plant responses are shared between conditions, while others are specific to either expression of P19 only (green), or co-expression of P19 and HA proteins (purple; see text for details). Black and grey arrows indicate strong or weak activation, respectively. Abbreviations: AsA, ascorbic acid; Chp, chloroplast; ET, ethylene; HA, hemagglutinin; HSP, heat shock protein; JA, jasmonic acid; JA-Ile, jasmonic acid isoleucine conjugate; SA, salicylic acid; SAR, systemic acquired resistance; VLP, virus-like particle.

Induced expression of *HSP* genes is common during heat shock stress, which interferes with general protein folding. P19-specific upregulation of these genes is thus consistent with high accumulation and localization of the VSR in the cytosol. Induced *HSP* gene expression is also a mark of SA signaling (Snyman and Cronjé, 2008; Sangwan *et al.,* 2022), consistent with enhanced SA accumulation and induction of SA response genes in P19 samples (Figure 7). Considering that *HSP* genes were not induced in H5 samples despite the co-expression of P19 (Figure 3), it is tempting to speculate that this response was somehow compromised by HA-induced oxylipin signaling, similar to how HA expression reduced SA accumulation and expression of SA response genes of the *PR1* and *PR2* families (Figure 7). Alternatively, P19 levels may be higher when the VSR is expressed alone, leading to higher SA responses.

Identification of HA-specific responses indicates that the secreted protein, or the accumulation of its associated VLPs, results in a unique molecular signature that reshapes the metabolism of plant cells. Among HA-induced genes, some encoded proteins involved in PA and LPA accumulation, or in the control of lipid distribution within membranes (Figure 4). Accumulation of PA likely contributes to immunity as this metabolite is a well-known messenger in stress signaling (Lim *et al.,* 2017). Alternatively, accumulation of specific lipids and altered distribution of membrane lipids may be consequences of the effects of the HA protein on the endomembrane system, including the budding of VLPs (Figure 12). Indeed, PA, DAG, and LPA exhibit conical shapes, and their localized accumulation promotes membrane bending rather than formation of planar bilayers (Roth, 2008). Enhanced accumulation or altered distribution of these lipids within membranes thus constitute key mechanisms to modulate membrane curvature. Interestingly, localized accumulation of conical-shaped lipids within the PM is exploited by the influenza virus during mammalian host cell entry and exiting via membrane budding (Oguin *et al.,* 2014). As for influenza virions, we hypothesize that modifications in lipid composition and distribution within membranes were due to feedback mechanisms caused by high production of secreted HAs and budding of VLPs.

Transcriptomics analyses also revealed that wounding and herbivory response genes were highly induced in HA-expressing samples (Figure 12). Quantification of jasmonates accordingly showed HA-specific accumulation of JA, but not JA-Ile (Figure 8B). In P19 and H5 samples, genes promoting accumulation of JA-Ile were correspondingly repressed (Figure 8E), suggesting that HA expression results in oxylipin-related responses that do not uniquely (or mainly) proceed through the JA pathway and that other bioactive oxylipins are produced in this condition (Figure 12). While these signaling compounds remain to be formally identified, oxidized polyunsaturated fatty acids, divinyl ethers, and cyclopentenones such as OPDA are likely candidates. In line with this, oxylipin regulatory genes from alternative branches of the octadecanoid pathway were strongly induced by HA protein expression (Figure 9), including 9*-LOX,* α*-DOX,* and *CYP74* genes. In tobacco, agroinfiltration was shown to induce 9-LOX activity and corresponding upregulation of *9-LOX* genes (Huang *et al.,* 2010). Likewise, our data shows that several oxylipin regulatory genes were induced in AGL1 and P19 samples, including *9-LOX* gene *NbLOX1* (Figure 9B). Following agroinfiltration, upregulation of these genes was however greatly enhanced by HA protein expression (Figure 12). In the same report (Huang *et al.,* 2010), 13-LOX and 13-hydroperoxide lyase (HPL) enzymes were shown to stimulate production of green leaf volatiles (GLVs) such as hexanal (Figure 9A). While the 13-LOX encoding gene *NbLOX3a* was induced in H5 samples, the *HPL* gene Niben101Scf00313g08016 (*NbHPL*) was strongly repressed in P19 and H5 samples (Figure 9B; Tables S1 and S9). This suggests that 13-LOX activity supports JA synthesis in H5 samples (Figure 8B), but that GLV production is not a major component of the plant response to VLPs.

In Arabidopsis and tomato, OPDA has signaling functions independent of JA or JA-Ile (Stintzi *et al.,* 2001; Bosch *et al.,* 2014). Genes specifically induced by OPDA were also identified in Arabidopsis (Taki *et al.,* 2005; Ribot *et al.,* 2008). In H5 samples, strong and specific upregulation of homologs from these OPDA-specific genes suggests that this signaling cascade is active during HA protein expression. These include *BBE* genes (Figure 5C), phosphate transporter gene Niben101Scf05712g02003 (*NbPHO1;H10*), zinc finger protein (ZFP) genes Niben101Scf05373g01004 (*NbZFP1*) and Niben101Scf10015g02003 (*NbZFP2*), as well as ribonuclease genes Niben101Scf04082g01004 (*NbRNS1*) and Niben101Scf09597g01002 (*NbRNS2*; Tables S1 and S9). For *ZFP* and ribonuclease genes, RTqPCR confirmed strong and H5-specific induction (Figure S3), further suggesting that oxylipins other than JA and JA-Ile are actively signaling in this condition (Figure 12).

While partially compromised in SA responses, H5 samples still displayed induction of SAR-related genes, including homologs of SAR regulatory gene *AZI1* (Figure 7H). As SAR is often linked to SA signaling, this result at first seemed contradictory. JA and other oxylipins however induce both local and systemic defenses (Vicente *et al.,* 2012; Bosch *et al.,* 2014), perhaps explaining the SAR signature in HA-expressing samples. Alternatively, and has shown in Arabidopsis, a subset of SAR-related genes not only responds to SA, but also to volatile terpenes that promote inter-plant communication in parallel with SA (Riedlmeier *et al.,* 2017). Within that subset, *AZI1*, *CHI*, *MLO6* and *BBE* genes can be found, a signature that resembles SAR-related genes induced by VLP expression (Figure 7). To investigate terpene signaling during foreign protein expression, RNAseq data was searched to identify terpene-related genes. This revealed upregulation of several candidates from the mevalonate pathway, which promotes assembly of terpene backbone molecules (Figure S4A). Also induced, were several *CYP71* genes that mediate diversification of terpene backbone molecules so that terpene blends with refined biological functions can be produced (Banerjee and Hamberger 2018). While induced expression of some terpene-related genes was seen in AGL1 and P19 samples, expression was either higher or specific to H5 samples (Figure S4B and Tables S1 and S12). As RTqPCR confirmed upregulation patterns of several terpene-related genes (Figure S4C), results suggest that terpene signaling is another key feature of the expression system (Figure 12), with VLP expression leading to stronger and broader activation of this pathway. In turn, stronger and broader terpene signaling from H5 samples may promote SAR despite compromised SA responses in this condition. In a molecular farming context where agroinfiltrated plants are packed in closed incubation chambers, plant-to-plant communication would be expected to play a role, especially since expression lasts for several days. Whether these responses are due to intra- or inter-plant signaling however remains to be determined, keeping in mind that every cell from infiltrated leaves and every leaf from infiltrated plants theoretically express VLPs.

Overall, our results highlight diversity and interconnectivity of the responses induced in *N. benthamiana* leaves expressing foreign proteins. While some of these responses will likely vary according to the foreign proteins expressed, data reported here is still of practical significance considering that *Agrobacterium*-mediated expression is widely employed for molecular farming. Obviously, this unique pathosystem remains artificial, yet it now contributes to solving major public health issues, including worldwide spreading of infectious diseases such as influenza. Despite stress imposed by the system, plants show remarkable resilience that allows them to produce high amounts of recombinant proteins. Our analyses will help to improve molecular farming techniques, as exemplified by the reduction of HA-induced defenses and stress symptoms by AsA (Figures 11 and 12). Although such treatments would be challenging to implement for commercial-scale production of VLPs, these results nonetheless demonstrate the importance of plant immunity during foreign protein expression. They also confirm that hypothesis-driven strategies can be applied to improve productivity, or as in this case biomass quality. Based on the knowledge developed, alternative improvement strategies can be envisioned, including targeted edition of the host plant genome and co-expression of protein helpers (Goulet *et al.,* 2012; Jutras *et al.,* 2015; Grosse-Holz *et al.,* 2018).

## Materials and methods

### Seed germination and plant growth

Seeds of *N. benthamiana* were spread on pre-wetted peat mix plugs (Ellepot) and placed in a germination chamber for 2 days, where conditions were as follows: 28°C/28°C day/night temperature, 16 h photoperiod, 90% relative humidity, and light intensity of 7 µmol m^-2^ s^-1^. Germinated plantlets were next transferred in a growth chamber for 15 days, where conditions were as follows: mean temperature of 28°C over 24 h, 16 h photoperiod, mean relative humidity of 66% over 24 h, 800 ppm carbon dioxide (CO_2_) injected only during the photo-phase, and light intensity of 150 µmol m^-2^ s^-1^. During this time, watering and fertilization were provided as needed. After 2 weeks, peat mix plugs were transferred to 4 inches pots containing pre-wetted peat-based soil mix (Agro-Mix). Freshly transferred plantlets were then moved to a greenhouse, where conditions were as follows: mean temperature of 25°C over 24 h, 16 h photoperiod, mean relative humidity of 66% over 24 h, 800 to 1,000 ppm CO_2_ injected only during the photo-phase, and light intensity according to natural conditions, but supplemented with artificial high pressure sodium lights at 160 µmol m^-2^ s^-1^. In the greenhouse, watering and fertilization were provided as needed. Growth was allowed to proceed for an average of 20 additional days, until the plants were ready for agroinfiltration.

### Binary vector constructs

For VLP expression, sequences from the mature HA protein of pandemic influenza virus strain H5 Indonesia (H5/A/Indonesia/05/2005; H5^Indo^) were fused to the signal peptide of a *Medicago sativa* (alfalfa) PDI using PCR-based methods. Once assembled, the chimeric *H5^Indo^* gene was reamplified by PCR and then introduced in the T-DNA region of a pCAMBIA binary vector previously linearized with restriction enzymes *SacII* and *StuI* using the In-Fusion cloning system (Clontech). Expression of *H5^Indo^* was driven by a 2X35S promoter from the cauliflower mosaic virus (CaMV). The expression cassette also comprised 5’- and 3’-untranslated regions (UTRs) from the cowpea mosaic virus (CPMV), and the *Agrobacterium nopaline synthase* (*NOS*) gene terminator. To prevent silencing induced by recombinant gene expression *in planta*, T-DNA region of the binary vector also included the suppressor of RNA silencing gene *P19*, under the control of a plastocyanin promoter and terminator. For P19 samples, a binary vector allowing expression of P19 only was employed. For AGL1 samples, a binary vector harboring a frameshifted version of *P19* was used as a control.

### Agrobacterium cultures and plant infiltration

Binary vectors were transformed by heat shock in *Agrobacterium* strain AGL1. Transformed bacteria were plated on Luria-Bertani (LB) medium, with appropriate antibiotics selection (kanamycin 50 µg/ml). Colonies were allowed to develop at 28°C for 2 days. Using isolated colonies, frozen glycerol stocks were prepared and placed at -80°C for long-term storage. When ready, frozen bacterial stocks were thawed at room temperature before transfer in pre-culture shake flasks containing LB medium with antibiotics selection (kanamycin 50 µg/ml). Bacterial pre-cultures were grown for 18 h at 28°C with shaking at 200 rpm. While keeping kanamycin selection, pre-cultures were transferred to larger shake flasks and bacteria were allowed to develop for an extra 18 h at 28°C with shaking at 200 rpm. Using a spectrophotometer (Implen), bacterial inoculums were prepared by diluting appropriate volumes of the bacterial cultures in resuspension buffer (10 mM MgCl_2_, 5 mM MES, pH 5.6). A final OD_600_ of 0.6 was employed for all experiments and vacuum infiltration was performed by submerging whole plant shoots in the appropriate bacterial suspension.

### Transient protein expression and biomass harvesting

Recombinant protein accumulation was allowed to proceed for 6 or 7 days, as indicated. For all experiments, expression took place in condition-controlled plant growth chambers, where settings were as follows: 20°C/20°C day/night temperature, 16 h photoperiod, 80% relative humidity, and light intensity of 150 µmol m^-2^ s^-1^. Watering was performed every other day, with no fertilizer supplied during the expression phase. For biomass harvesting, leaves of similar developmental stage were selected using the leaf plastochron index (Meicenheimer, 2014). The fourth and fifth fully expanded leaves starting from the top of each plant were harvested without petiole. Freshly cut leaves were placed in pre-frozen 50 mL Falcon tubes, before flash freezing in liquid nitrogen. Frozen biomass was stored at -80°C until ready for analysis. Using pre-chilled mortars and pestles, foliar tissue was ground and homogenized into powder using liquid nitrogen. Each sample was made from four leaves collected on two randomly selected plants. The average results presented were obtained from at least three biological replicates.

### Protein extraction, western blotting, and HMG assays

For protein extraction, 1 g of frozen biomass powder was taken out of the -80°C freezer and placed on ice. A 2 mL volume of extraction buffer (50 mM Tris, 500 mM NaCl, pH 8.0) was added, followed by 20 µL of 100 mM phenylmethanesulfonyl fluoride (PMSF) and 2 µL of 0.4 g/mL metabisulfite. Quickly after addition of all solutions, samples were crushed for 45 sec using a Polytron homogenizer (ULTRA-TURRAX^®^ T25 basic) at maximum speed. One mL of each sample was transferred to a prechilled eppendorf tube and centrifuged at 10 000 x g for 10 min at 4°C. Supernatants were carefully recovered, transferred to new eppendorf tubes, and kept on ice until determination of protein concentration. To quantify protein content from crude extracts, the Bradford method was employed, with bovine serum albumin as a protein standard.

For western blotting, total protein extracts were diluted in extraction buffer and mixed with 5X Laemmli sample loading buffer to reach a final concentration of 0,5 µg/µL. Protein samples were denatured at 95°C for 5 min, followed by a quick spin using a microcentrifuge. 20 µL of each denatured protein extract (10 µg) was then loaded on Criterion^TM^ XT Precast polyacrylamide gels 4-12% Bis-Tris and separated at 110 volts for 105 min. Using transfer buffer (25 mM Tris, 192 mM Glycine, 10% methanol), proteins were next electrotransferred onto a polyvinylidene difluoride (PVDF) membrane at 100 volts. After 30 min, membranes were placed in blocking solution: 1X Tris-Buffered Saline with Tween-20 (TBS-T; 50 mM Tris, pH 7.5, 150 mM NaCl, 0,1% (v/v) Tween-20), with 5% nonfat dried milk. Membranes were blocked overnight at 4°C with gentle shaking. The next morning, blocking solution was removed and primary antibodies were incubated at room temperature for 60 min with gentle shaking in 1X TBS-T, 2% nonfat dried milk solution. After 4 washes in 1X TBS-T, secondary antibodies were added and incubated at room temperature for 60 min with gentle shaking in 1X TBS-T, 2% nonfat dried milk solution. After 4 extra washes in 1X TBS-T, Luminata^TM^ Western HRP Chemiluminescence Substrate (Thermo Fisher Scientific) was added to the membranes and protein complexes were visualized under the chemiluminescence mode of an Imager 600 apparatus (Amersham). Antibody dilutions were as follows: anti-HA A/Indonesia/05/2005 (H5N1; CBER): 1/5,000 (primary antibody). Rabbit anti-sheep (JIR): 1/10,000 (secondary antibody).

For HMG assays, turkey red blood cells were diluted to a concentration of 0,25% (v/v) in phosphate-buffered saline solution (PBS; 0.1 M PO_4_, 0.15 M NaCl, pH 7.2). While keeping red blood cells on ice, protein samples were diluted in extraction buffer using 1/384 and 1/576 ratios. For each dilution, 200 µL of total protein extract was transferred to the first row of a 96-well plate. Eight serial dilutions were then performed using 100 µL of protein extract mixed to 100 µL of PBS buffer previously poured in each plate well. Following serial dilutions, 100 µL of red blood cell solution was added to protein extracts. After thorough mixing, samples were incubated overnight at room temperature. HA activity was scored visually on the next day.

### RNA extractions and quantification

Using the RNeasy commercial kit (Qiagen), 100 mg of frozen biomass powder was used for RNA extractions. Residual DNA was removed using the RNase-free DNase Set (Qiagen). Concentration of RNA extracts was determined using a spectrophotometer (Implen) and integrity evaluated using a 2100 BioAnalyzer (Agilent). For long term storage, RNA extracts were stabilized by adding the RNAseOUT recombinant ribonuclease inhibitor (Thermo Fisher Scientific), before freezing at -80°C until ready for further analysis.

### RTqPCR analyses

For each sample, 1 µg of RNA was reverse transcribed into cDNA using the QuantiTect Reverse Transcription Kit (Qiagen). Transcript quantification was performed in 96-well plates, using the ABI PRISM 7500 Fast real-time PCR system and custom data analysis software (Thermo Fisher Scientific). Each reaction contained the equivalent of 5 ng cDNA as a template, 0.5 µM of forward and reverse primers, and 1X QuantiTect SYBR Green Master Mix (Qiagen) for a total reaction volume of 10 µL. RTqPCR runs were done under the SYBR Green amplification mode and cycling conditions were as follows: 15 min incubation at 95°C, followed by 40 amplification cycles at 95°C for 5 sec, 60°C for 30 sec, and 65°C for 90 sec. Reactions in the absence of cDNA template were conducted as negative controls and melting curve analyses were performed to confirm the lack of primer dimer formation and amplification specificity. Resulting fluorescence and cycle threshold (Ct) values were next exported to the Microsoft Excel software. To correct for biological variability and technical variations during RNA extraction, quantification, or reverse transcription, expression from six housekeeping genes (*NbACT1*, *NbVATP*, *NbSAND*, *NbUBQ1*, *NbEF1-*α, and *NbGAPDH1*) was used to normalize expression data (Vandesompele *et al.,* 2002). Stable expression of housekeeping genes was verified by RTqPCR (Figure S5) and using RNAseq data (Table S13). Normalized numbers of molecules per ng of RNA were deduced using the 2^-ΔΔCt^ method (Livak and Schmittgen, 2001; Bustin *et al.,* 2009) and standard curves derived from known quantities of phage lambda DNA. Standard deviation related to the within-treatment biological variation was calculated in accordance with the error propagation rules and sequences of all primers used in this study are available in the Table S14.

### RNAseq analyses

To study global changes to the plant transcriptome, RNA extracts were placed on dried ice and shipped to an external service provider (Génome Québec; McGill University, Montréal). Briefly, ribosomal RNA (rRNA) was depleted to concentrate messenger RNA (mRNA). After fragmentation, mRNA stranded nucleotide libraries were created by fusing TruSeq RNA sequencing adapters (Illumina), followed by library amplification by PCR. High throughput sequencing was performed using a HiSeq 2000 SR100 device (Illumina). Three biological replicates from each experimental condition were sequenced. For each library, an average of 36 954 631 reads were obtained and a total of 3 695 463 144 bases were sequenced. Following filtering and quality control steps, sequences were aligned against the nuclear genome of *N. benthamiana* (https://solgenomics.net/). Once aligned and quantified, sequencing data was used to perform pairwise comparisons between conditions, as indicated.

### Large-scale proteomics

To study changes in protein abundance, total protein extracts from three biological replicates of each condition were used for iTRAQ labeling. Detailed description of sample preparation, peptide labeling, and mass spectrometry methods are available elsewhere (Jutras *et al.,* 2020). Peptide mass spectra were analyzed using the Mascot database search engine (Matrix Science), which is implemented in Proteome Discoverer (Thermo Scientific). The search algorithm was set to interrogate the Solanaceae protein database from UniProt, which comprises over 117 800 proteins (http://www.uniprot.org/taxonomy/4070). Common peptide contaminants like keratin were taken out of the analysis.

### Quantification of stress hormones

For quantification of salicylates and jasmonates, 500 mg of fresh leaf tissue was employed. For quantification of ACC, 1 g of frozen leaf biomass powder was lyophilized for 48 h to obtain 50 mg of dried tissue. Once weighted, samples were placed on dry ice and shipped to Dr. Irina Zaharia at the National Research Council of Canada located in Saskatoon, Saskatchewan, Canada. Hormone quantification was performed using high-performance liquid chromatography-electrospray ionization-tandem mass spectrometry (HPLC-ESI-MS/MS), using the ACQUITY UPLC system equipped with a binary solvent delivery manager and a sample manager (Waters). This system was also coupled to a Micromass Quattro Premier XE quadrupole tandem mass spectrometer (Waters). MassLynx and QuanLynx softwares (Micromass) were used for data acquisition and analysis. Quantification data was obtained from comparison with standard-derived calibration curves.

### Lignin quantification

Lignin quantification was performed using an acid-catalyzed reaction that forms soluble lignin-thioglycolate complexes. Suitable for photometric measurements, this method was developed in spruce (*Picea abies*; Lange *et al.,* 1995), but later adapted for use in *N. benthamiana* (de la Torre *et al.,* 2014). For each sample, 200 mg of grounded biomass was used. Quantification of lignin was obtained by measuring absorbance at 280 nm and comparison with standard curves made using known quantities of purified lignin polymers (Sigma).

### Ascorbic acid treatments

Purified AsA was purchased from Sigma. A 10 mM solution was obtained by dilution in 20 mM MES (pH 6.0), supplemented with 0.05% Tween 20. After solubilization of AsA, the pH was readjusted to 6.0 using 5 N NaOH, before spraying on the plants. For each treatment, blocks of 12 plants were uniformly sprayed with a 50 mL volume of the AsA solution, or with a similar volume of the mock solution at pH 6.0. A first treatment was applied at 2 DPI. Recall treatments were then performed at 4 and 6 DPI, before harvesting at 7 DPI. Untreated NI, P19, and H5 plants were used as controls.

### Statistical analyses

Statistical analyses were performed on Graph Pad Prism 9.5.0. Groups were analysed using one-way ANOVA followed by a post-hoc Tukey’s multiple comparison test with an alpha threshold of 0.05. Groups are labeled with a compact letter display. Groups not sharing the same letter are statistically different.

### Accession numbers

Raw sequencing data from the RNAseq study is available on the Gene Expression Omnibus (GEO) website under the accession number GSE233178.

## Acknowledgments and funding

This work would not have been possible without support from the Biomass Production and Research & Innovation teams of Medicago. Their contribution includes but is not limited to cloning of the genetic constructs, inoculum preparation, growth of the plants and agroinfiltration. Authors also wish to acknowledge Gervais Pelletier and Dr. Armand Séguin from Natural Resources Canada, for technical support and advice on RTqPCR. For support during the proteomics study, authors are also grateful to Antoine Leuthreau, graduate student at the Department of phytology from Laval University. Authors finally wish to thank Dr. Sébastien Mongrand for helpful discussion on plant lipids. Funding for this work was provided by Medicago Inc. This work was also supported by Collaborative Research and Development grants from the Natural Sciences and Engineering Research Council of Canada to Dominique Michaud (CRDPJ 495852) and Peter Moffett (CRDPJ 477619).

## Author contributions

L.P.H., D.M., P.M., and M.A.D. designed the research, supervised the project, and analyzed the data. L.P.H. and M.A.C. drafted the manuscript and assembled the figures. P.O.L. managed production of genetic constructs. R.T. performed RTqPCR, HMG assays and western blots. F.P.G. and M.A.C. performed statistical analyses. L.P.H., A.R. G.G, and C.B. performed RNAseq. J.F.L., M.A.C., and F.P.G. generated RNAseq *in silico* data. M.C.G., A.B., and L.P.H. conducted proteomics studies and produced biomass for plant hormone quantification. M.C.G and F.P.G. generated proteomics *in silico* data. E.R. performed lignin quantification. R.T. and L.P.H. performed experiments with AsA. All authors read, helped to edit, and approved final version of the manuscript.

## Data availability statement

All data discussed in this study can be found in the manuscript and in the Supplementary Materials.

## Supporting information

Additional supporting information may be found online in the Supporting Information section at the end of the article.

**Table S1.** Unsorted lists of genes with significantly altered expression in AGL1, P19, and H5 samples.

**Table S2.** Lists of chloroplast-related genes found in P19 and H5 samples.

**Table S3.** Lists of *HSP* and chaperone genes found in P19 and H5 samples.

**Table S4.** List of lipid-related genes found in P19 and H5 samples.

**Table S5.** List of genes related to oxidative stress activation found in P19 and H5 samples.

**Table S6.** List of genes related to sugar metabolism found in P19 and H5 samples.

**Table S7.** List of lignin-related genes found in P19 and H5 samples.

**Table S8.** List of genes related to SA signaling and SAR found in P19 and H5 samples.

**Table S9.** Lists of oxylipin-related genes found in P19 and H5 samples.

**Table S10.** List of ET- and senescence-related genes found in P19 and H5 samples.

**Table S11.** Unsorted lists of proteins with significantly altered accumulation levels in AGL1, P19, and H5 samples.

**Table S12.** List of terpene-related genes found in P19 and H5 samples.

**Table S13.** RNAseq data from reference genes used to normalize RTqPCR results.

**Table S14.** List of RTqPCR primers used in this study.

**Figure S1.**
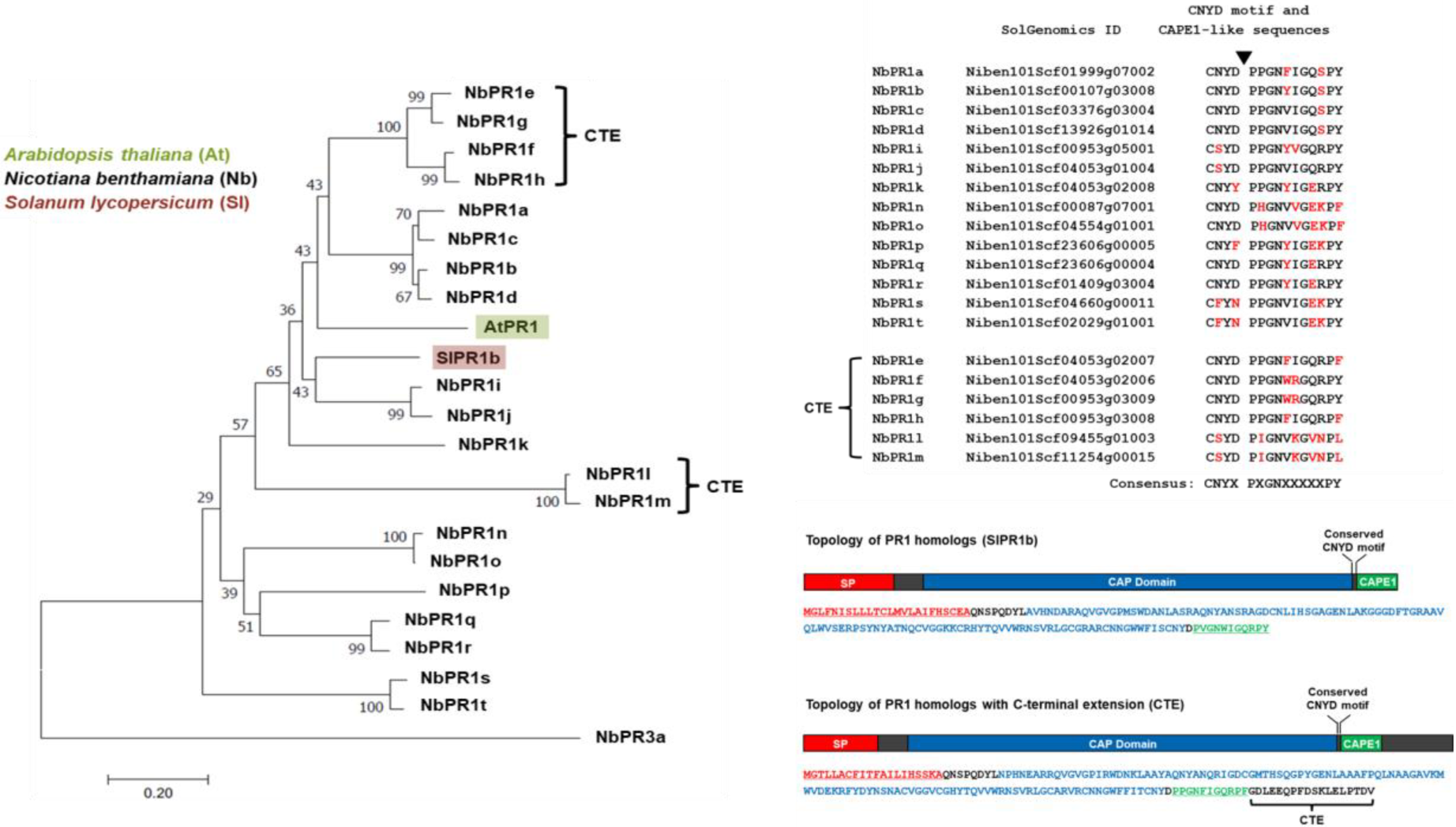
Phylogenetic relationships between PR1 homologs from *N. benthamiana*. The genome of *N. benthamiana* was searched using full-length amino acid sequences of *Solanum lycopersicum* (tomato) SlPR1b (accession number NP_001234314) or Arabidopsis AtPR1 (At2g14610) as queries. Full-length protein sequences that were retrieved were next aligned with ClustalW, using chitinase NbPR3a (Niben101Scf07491g00003) as an outgroup. Alignment parameters were as follow: for pairwise alignment, gap opening, 10.0, and gap extension, 0.1; for multiple alignment, gap opening, 10.0, and gap extension, 0.20. Resulting alignments were submitted to the MEGA5 software and a neighbor-joining tree derived from 5000 replicates was generated. Bootstrap values are indicated on the node of each branch. At the top right, gene model number from corresponding *NbPR1* genes are shown, along with amino acid sequences of the conserved CNYD motif and CAPE1 peptide. A black arrow indicates clipping site of the CAPE1 peptide. At the bottom right, topologies from PR1 proteins with or without a C-terminal extension (CTE) are shown. Abbreviations: At, *Arabidopsis thaliana*; Nb, *Nicotiana benthamiana*; Sl, *Solanum lycopersicum*; SP, signal peptide.

**Figure S2.**
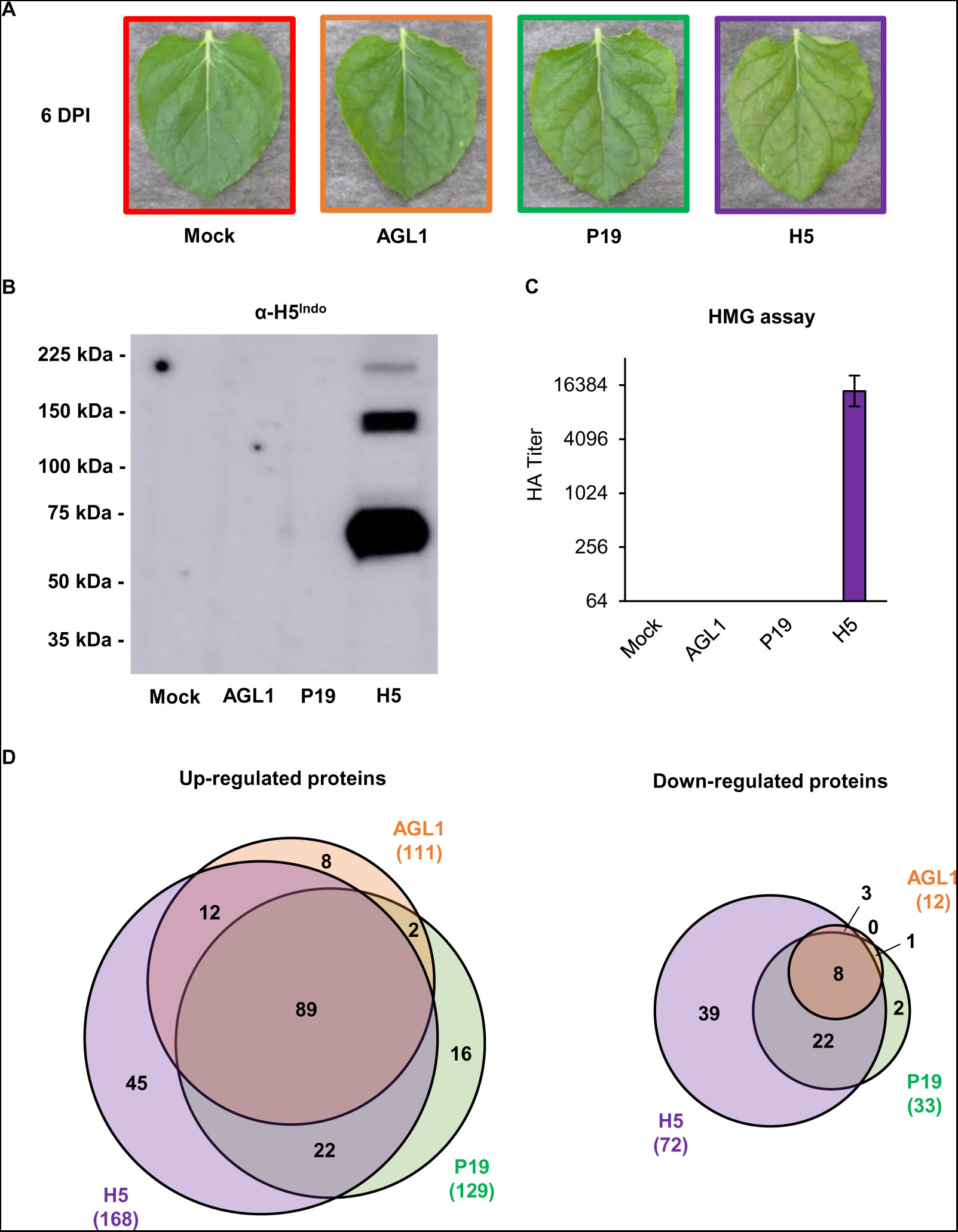
Stress symptoms and analysis of biomass used for proteomics. (A) Stress symptoms observed on representative leaves from each condition harvested at 6 DPI. (B) Western blot confirming HA protein accumulation. (C) Hemagglutination (HMG) assay confirming HA protein activity. (D) Venn diagrams depicting up- and downregulated proteins from pairwise comparisons: AGL1 *vs* Mock, P19 *vs* Mock, and H5 *vs* Mock. Circle size is proportional to the number of proteins significantly regulated. Proteins specific to AGL1 infiltration are shown in orange. Proteins specific to P19 expression are shown in green. Proteins specific to P19 and HA co-expression are shown in purple. Diagram intersects show proteins common to more than one condition. Condition names are as follow: Mock: leaves infiltrated with buffer only; AGL1: leaves infiltrated with *Agrobacterium* strain AGL1 that carry a binary vector control; P19: leaves infiltrated with AGL1 and expressing P19 only; H5: leaves infiltrated with AGL1 and co-expressing P19 and H5^Indo^.

**Figure S3.**
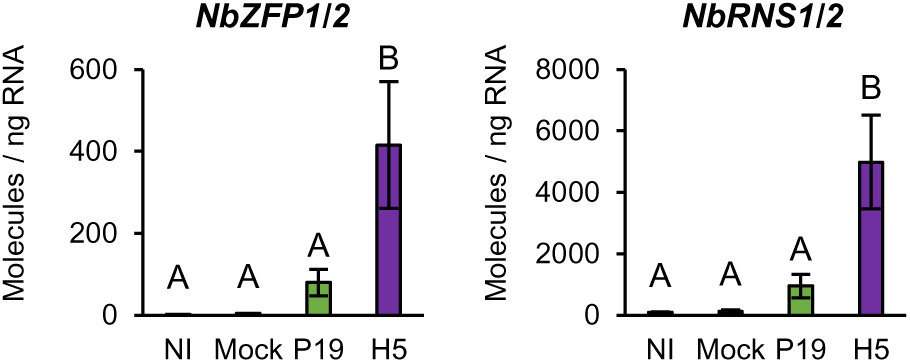
Expression of OPDA-specific genes. Expression of genes specifically induced by OPDA as measured by RTqPCR at 6 DPI. Results are expressed in numbers of molecules per ng of RNA. Groups that do not share the same letter are statistically different. Condition names are as follow: NI: non-infiltrated leaves; Mock: leaves infiltrated with buffer only; P19: leaves infiltrated with AGL1 and expressing P19 only; H5: leaves infiltrated with AGL1 and co-expressing P19 and H5^Indo^.

**Figure S4.**
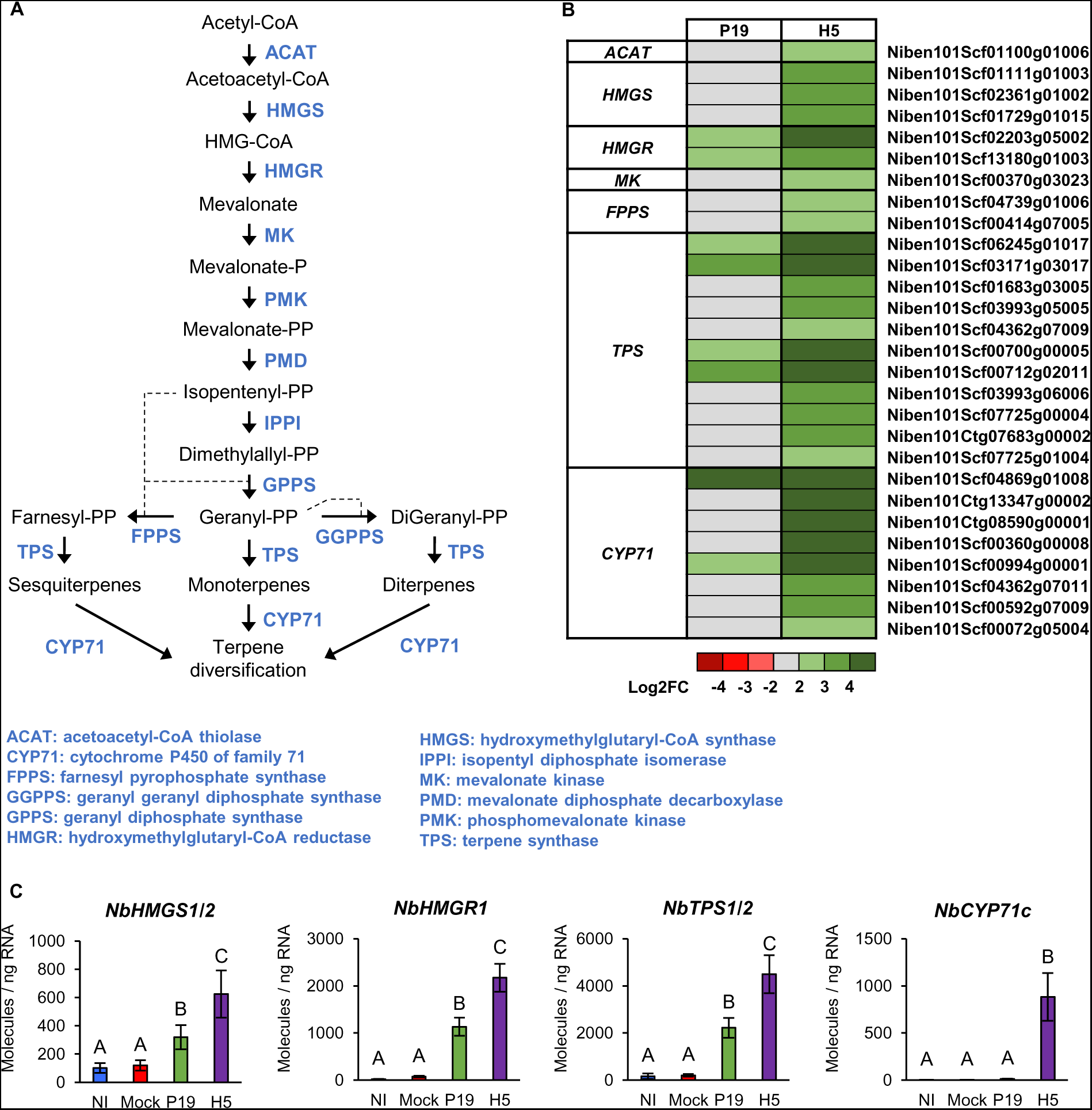
Expression of terpene-related genes. (A) Overview of the mevalonate pathway. Metabolic intermediates are shown in black while enzymes are shown in blue. (B) Heat-map depicting expression of genes involved in terpene synthesis and diversification at 6 DPI. Each line represents a gene shown in the Table S12. Grey indicates genes that are not differentially expressed. Green are red colored gradients respectively reflect extent of gene up- and downregulation, as indicated. (C) Expression of genes involved in terpene synthesis and diversification as measured by RTqPCR at 6 DPI. Results are expressed in numbers of molecules per ng of RNA. Groups that do not share the same letter are statistically different. Condition names are as follow: NI: non-infiltrated leaves; Mock: leaves infiltrated with buffer only; P19: leaves infiltrated with AGL1 and expressing P19 only; H5: leaves infiltrated with AGL1 and co-expressing P19 and H5^Indo^.

**Figure S5.**
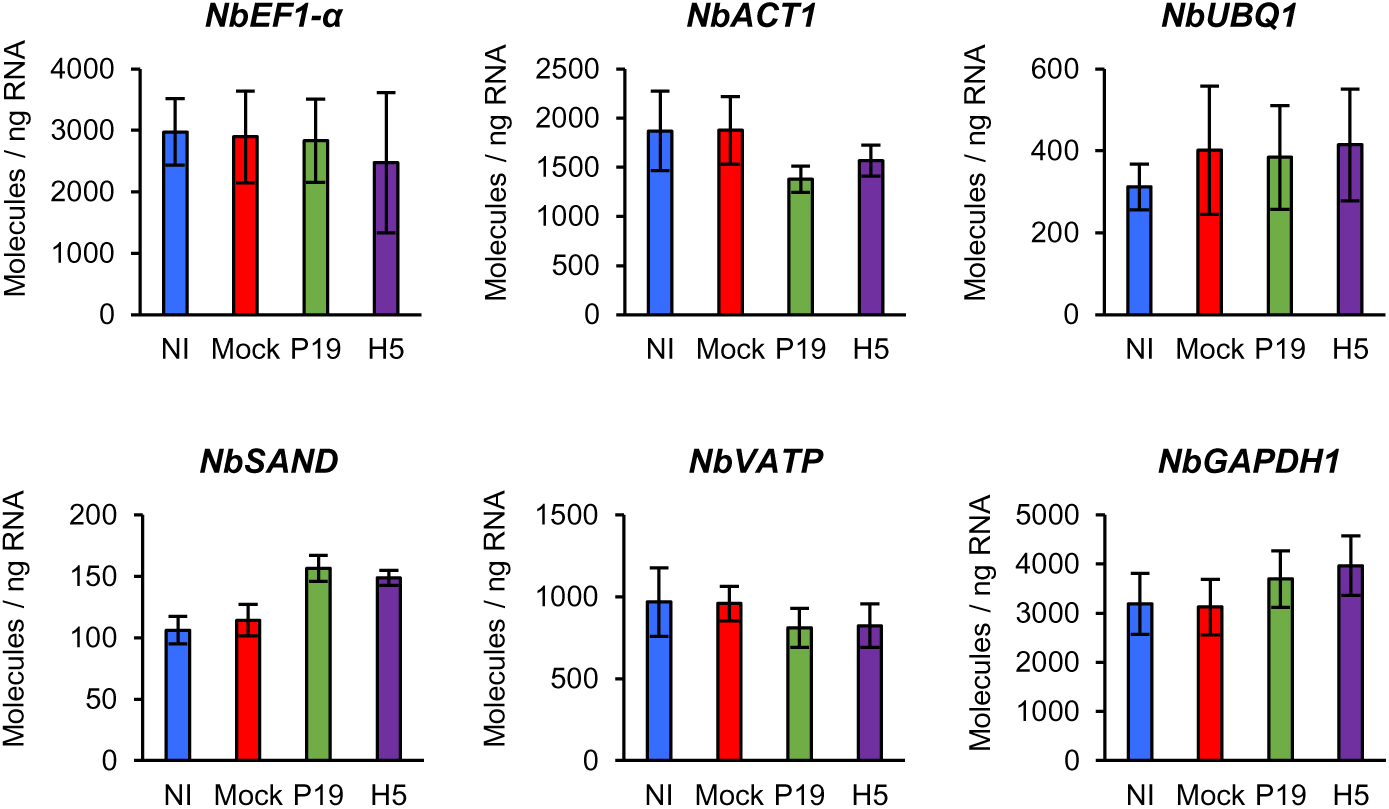
Expression of reference genes. Expression of housekeeping genes used to normalize RTqPCR data at 6 DPI. Results are expressed in numbers of molecules per ng of RNA. No statistically significant difference was observed between the groups. Condition names are as follow: NI: non-infiltrated leaves; Mock: leaves infiltrated with buffer only; P19: leaves infiltrated with AGL1 and expressing P19 only; H5: leaves infiltrated with AGL1 and co-expressing P19 as well as H5^Indo^.

